# Co-occurring ripple oscillations facilitate neuronal interactions between cortical locations in humans

**DOI:** 10.1101/2023.05.20.541588

**Authors:** Ilya A. Verzhbinsky, Daniel B. Rubin, Sophie Kajfez, Yiting Bu, Jessica N. Kelemen, Anastasia Kapitonava, Ziv M. Williams, Leigh R. Hochberg, Sydney S. Cash, Eric Halgren

**Affiliations:** Neurosciences Graduate Program, University of California San Diego, La Jolla, CA 92093, USA; Medical Scientist Training Program, University of California San Diego, La Jolla, CA 92093, USA; Center for Neurotechnology and Neurorecovery, Department of Neurology, Massachusetts General Hospital, Boston, MA 02114, USA; Harvard Medical School, Boston, MA 02114, USA; Department of Radiology, University of California San Diego, La Jolla, CA 92093, USA; Department of Neurosciences, University of California San Diego, La Jolla, CA 92093, USA; Department of Neurosurgery, Massachusetts General Hospital, Boston, MA 02114; Program in Neuroscience, Harvard-MIT Program in Health Sciences and Technology, Harvard Medical School, Boston, MA 02115; Center for Neurorestoration and Neurotechnology, Department of Veterans Affairs, Providence, RI 02908, USA; Carney Institute for Brain Science and School of Engineering, Brown University, Providence, RI 02912, USA

## Abstract

Synchronous bursts of high frequency oscillations (‘ripples’) are hypothesized to contribute to binding by facilitating integration of neuronal firing across cortical locations. We tested this hypothesis using local field-potentials and single-unit firing from four 96-channel microelectrode arrays in supragranular cortex of 3 patients. Neurons in co-rippling locations showed increased short-latency co-firing, prediction of each-other’s firing, and co-participation in neural assemblies. Effects were similar for putative pyramidal and interneurons, during NREM sleep and waking, in temporal and Rolandic cortices, and at distances up to 16mm. Increased co-prediction during co-ripples was maintained when firing-rate changes were equated, and were strongly modulated by ripple phase. Co-ripple enhanced prediction is reciprocal, synergistic with local upstates, and further enhanced when multiple sites co-ripple. Together, these results support the hypothesis that trans-cortical co-ripples increase the integration of neuronal firing of neurons in different cortical locations, and do so in part through phase-modulation rather than unstructured activation.

## INTRODUCTION

The ‘Binding Problem’ describes a fundamental question in systems neuroscience: how are the different elements of a mental event unified into a cohesive experience despite being encoded in locations distributed across the cortex? The mechanisms supporting cortical binding are poorly understood, but one proposed mechanism, ‘binding-by-synchrony’ (BBS), posits that high frequency oscillations, synchronized between widespread cortical areas, form transient integrated networks of activity across the cortex^1,2^. In this model, rhythmic pulses of depolarization in different cortical locations modulate neuronal firing such that their cells fire in coordinated spatial patterns. Considerable evidence supporting^3^ and questioning^4,5^ this hypothesis has been obtained, but studies have been mainly confined to the visual system of rodents and cats, and whether it is tenable in the human neocortex is unclear.

In a seemingly unrelated stream of research, high frequency oscillations (‘ripples’) in rodent hippocampus during non-rapid eye movement (NREM) sleep have been shown to organize the firing of cells, replaying those encoding events from previous waking periods, critical for memory consolidation in the cortex^6-9^. Ripples also occur in rodent association cortex, where they couple with hippocampal ripples during sleep following spatial memory tasks^10^.

Similar events have recently been found in humans during waking and NREM sleep^11-17^. Hippocampal formation and cortical ripple occurrence and co-occurrence increase preceding memory recall^11,12,15^, and cortical neuron firing sequences established during encoding replay during ripples prior to recall^18^ and during resting or NREM sleep when consolidation may occur^19,20^. These findings are consistent with cortical ripples contributing to memory consolidation and recall in humans. Specifically, ripple co-occurrence could facilitate the binding of different elements of memories that are represented in disparate cortical areas, the essence of hippocampus-dependent memory^21^.

Recently, we found that ∼100ms long ∼90Hz ripples are ubiquitous in all regions of the cortex during NREM as well as waking^14^. During waking, cortical ripples occur on local high frequency activity peaks. During sleep, cortical ripples occur, often during spindles, and typically on the down-to-upstate transition, with unit-firing patterns consistent with generation by pyramidal-interneuron feedback^14,22^. Ripples co-occur, and remarkably, phase-synchronize across all lobes and between both hemispheres, with little decrement, even at long distances^15^.

Ripples’ widespread co-occurrence and synchrony suggest they may serve a role in binding neural activity across disparate regions of the brain that is more general than their putative involvement in memory consolidation and recall. Studies of intracranial microelectrodes implanted in humans have shown that cortical ripples modulate local neuronal firing in a phase dependent manner^12,14,23^. However, a core prediction of the BBS hypothesis extends beyond local firing and posits that gamma-band synchrony across multiple sites facilitates integration of cortical computation across locations through phase selection of local neural activity^2^. This concept proposes that co-occurring high frequency oscillations comodulate involved neurons, enhancing their pairwise interactions and their impact on downstream circuits. While ripples have been shown to synchronize across cortical locations, the impact of co-rippling on the integration of firing across these areas has not been tested in humans.

Here, we analyzed a rare collection of intracranial Utah Array (UA) recordings with 96 fine tipped microelectrodes at 400 μm pitch implanted into cortical supragranular layers II/III, each spanning a ∼16 mm^2^ region of the cortex. We include multi-hour recordings from single arrays implanted in the temporal lobe of two patients undergoing evaluation of pharmaco-resistant epilepsy, and dual arrays simultaneously implanted in the primary motor cortex of a patient with tetraplegia participating in the BrainGate clinical trial^20,24^. We detected and classified putative pyramidal (PY) and interneuron (IN) units along with ripple oscillations in the local field potential (LFP) in NREM sleep and spontaneous waking. We show that co-occurring ripples increase co-firing between cells up to 16 mm apart. Co-ripples organize firing latencies and increase the ability of neurons to predict the activity of other neurons in other cortical locations. The increase in pairwise neural prediction is modulated by the relationship between the unit and ripple phase for neurons in both the target and predicting sites. Finally, we show that the expression of neural assemblies formed by groups of neurons is increased when member neurons are engaged in co-ripples. Together, these results support the fundamental core of the BBS hypothesis, that co-occurring ripple oscillations increase the integration of neuronal firing of neurons in different cortical locations, and do so in part through phase-modulation.

## RESULTS

| Array Implantation |  |  |  |  |  |  | Units |  |
| --- | --- | --- | --- | --- | --- | --- | --- | --- |
| Patient | Age | Sex | Reason for implant | Location | NREM duration (min) | Waking duration (min) | Total PY | Total IN |
| E1 | 51 | F | Epilepsy | Left anterior Middle Temporal Gyrus | 340 | 154 | 138 | 29 |
| E2 | 45 | M |  | Right anterior Superior Temporal Gyrus | 218 | N/A | 179 | 16 |
| B1 - dorsal | 36 | M | Spinal Cord Injury | Dorsal primary motor cortex | 695 | 173 | 79 | 15 |
| B1 - ventral |  |  |  | Ventral primary motor cortex |  | N/A | 41 | 12 |

### Characterization of units and ripples in motor cortex and temporal lobe

We analyzed Utah Array recordings (Fig. 1A) from two patient groups (Fig. 1B). Patients E1 and E2 had arrays implanted on the surface of the anterior temporal gyrus. Clinical electrodes were implanted in these patients for ∼7d to define the margins of the resection planned to treat their medication-resistant focal epilepsy. The clinical team determined prior to implantation that there was a very high probability that the resection would include the location where the Utah Array was placed, and this was indeed the case. Patient B1 suffered from tetraplegia secondary to a cervical spinal cord injury. Two arrays were implanted in the primary motor cortex of patient B1 as part of the BrainGate clinical trial to develop a brain-controlled movement prosthesis^20^. The cortex at the electrode sites is thought to be healthy, but the patient did have encephalomalacia in the left inferior frontal lobe. All patients gave their fully informed consent, and all procedures were monitored and approved by the responsible human subject protection committees. 418 ± 248 min of NREM was selected from all arrays and 154, 173 min of spontaneous waking was selected from patient E1 and the dorsal array in patient B1 (Table 1, Figure 1F,G). A total of 437 PY and 72 IN were sorted and classified from 254 microelectrode channels (127 ± 65 units per array, Fig. 1C,D). Ripples were detected during NREM and waking based on previously described methods^14,15^. Ripples per-channel density, amplitude, duration, and frequency was 18.6 ± 6.0 min^-1^, 14.2 ± 5.3 μV, 107 ± 19 ms, 89.3 ± 0.9 Hz during NREM and 12.4 ± 4.7 min^-1^, 18.6 ± 4.9 μV, 97 ± 29 ms, 89.8 ± 0.9 Hz during waking. Ripple characteristics are similar to those previously found^12,14,15^ and did not substantially differ between seizure patients and the BrainGate patient (See Supplementary Fig. 2). When a ripple detected on the same microelectrode contact contained a spike, there was an average of 1.63 ± 0.60 PY spikes and 2.08 ± 1.14 IN spikes during the single ripple event in NREM and 1.58 ± 0.47 PY spikes and 1.72 ± 0.86 IN spikes in waking. No comparisons were made between units recorded on the same channel.

**Figure 1.**
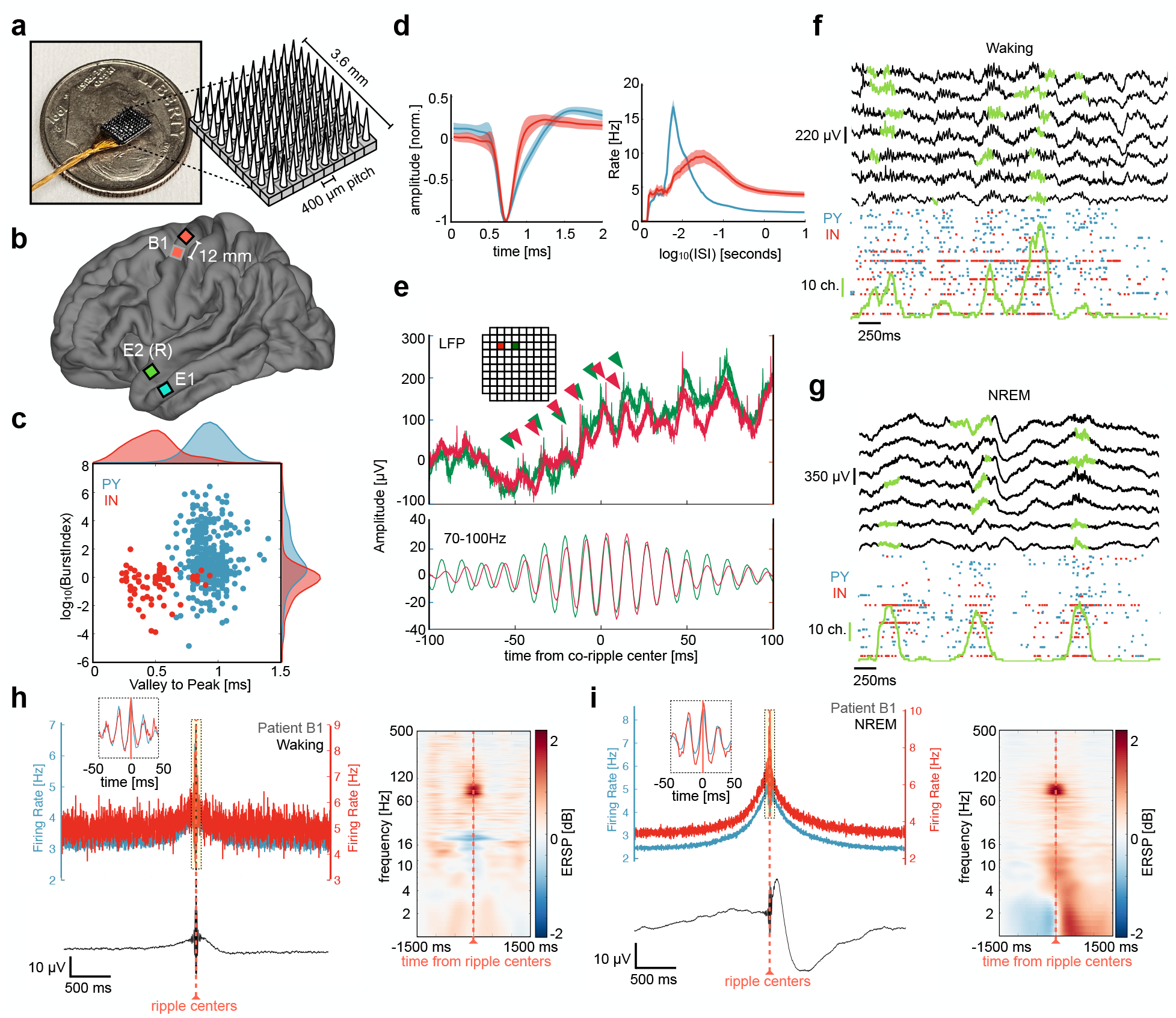
Utah Array implantation, unit firing during detected ripples. **a** Image of microelectrode configuration across array. **b** Array implantation locations for all patients (2 epilepsy, 1 BrainGate with 2 arrays). **c** Burst index and valley-to-peak time shown for putative PY (blue) and IN (red) units detected across all arrays. **d** Left, mean ± standard deviation waveforms shown for PY and IN. Right, mean ± SEM inter-spike interval (ISI) plots shown for PY and IN. **e** Top, example of broadband 30 kHz LFP data for a co-ripple occurring across two contacts (red and green). Carrots show single unit action potentials occurring on ripple peaks for each contact. Inset show the location of each contact on Utah Array (separated by 0.4 mm). Bottom, 70-100 Hz filtered data showing phase synchrony in the ripple band. **f** Top, sweep of broadband (0.1 – 1000Hz) LFP in 7 example microelectrodes during waking with detected ripples in green. Bottom, raster plot of associated PY and IN spiking with the number of microelectrodes containing a detected ripple in green (vertical green scale bar represents 10 microelectrode channels). **h** Top left, PY and IN firing rates locked to local ripple centers during waking. Bottom left, mean broadband LFP locked to ripple centers. Right, average time-frequency plots locked to microelectrode ripple centers. **g, i** same as in **f, h** but during NREM.

#### Unit firing increases during and phase locks to ripples in motor cortex and temporal lobe

Before testing how co-occurring ripples modulate unit co-firing, we measured how unit firing rates changed during ripples. Spike rates for each unit were quantified in motor cortex and temporal lobe during ripples detected on the unit’s channel and compared to baseline periods, which were randomly selected epochs in between ripples on the same channel. Both PYs and INs increased firing during ripples in the motor cortex and temporal lobe during NREM and waking. The mean ± st. dev. baseline spike rate for PYs and INs was 1.43 ± 1.81 Hz, and 2.82 ± 3.65 Hz in NREM, and 3.30 ± 2.52 Hz and 4.70 ± 6.52 Hz in waking. During ripples, the spike rates for PYs and INs increased to 4.19 ± 5.25 Hz and 8.46 ± 11.34 Hz in NREM, and 5.76 ± 5.52 Hz and 8.85 ± 11.37 Hz in waking (p_PY_NREM_ = 7.05e-36, p_IN_NREM_ = 3.75e-10, p_PY_wake_ = 6.08e-17, p_IN_wake_ = 3.22e-6, one sample two-sided Wilcoxon signed-rank).

In addition to increasing firing rates during the ripple period, our data^14^ and others^12,23^ have shown that firing tends to oscillate with ripple phase (Fig 1H,I). We quantified the extent to which the neurons in the data included in this study phase-locked with ripples. 272/437 (62%) PY and 61/72 (85%) IN phase-locked to ripples in NREM, while 147/217 (68%) PY and 35/44 (80%) IN phase-locked to ripples in waking (binomial test of spike ripple phase distributions within 0 ± π/2 vs. π ± π/2, expected value = 0.5, false discovery rate (FDR)^25^ corrected p < 0.05).

### Ripples co-occur within and across arrays with zero phase lag

Co-ripple rate (i.e., the percent of ripples on two contacts that overlap by 25ms or more) was greatest for adjacent contacts in the temporal lobe, falling rapidly and reaching a plateau of 15% at a distance of ∼1.5 mm (Fig 2A). The motor cortex demonstrated comparable overall co-ripple rates but decayed across a wider spatial scale (Fig 2B). The BBS hypothesis posits that high frequency oscillations not only co-occur, but phase synchronize across disparate brain regions. Therefore, in addition to co-occurrence rate, we evaluated phase synchrony across the array during co-ripples using the phase-locking value (PLV), a measure of phase-lag consistency invariant to amplitude^26^. The PLV during co-ripples was elevated across all distances compared to NREM epochs absent of detected ripples (Bonferroni-corrected α = 0.001 for 10 distance bins, p ≈ 0). Similar to co-occurrence rate, PLV decayed to a plateau in the temporal lobe, while remaining constant up to 4mm in the motor cortex (Fig. 2C,D). Given that the boundaries of phase synchronized modules were visible within arrays in the temporal lobe, we next estimated their size using non-negative matrix factorization (NMF), a location-naїve clustering algorithm. In the temporal lobe, the mean NMF cluster size was 1.29 ± 0.12 mm during NREM and 1.21 ± 0.09 mm during waking, which is consistent with previous reports of functional modules in the temporal lobe^27^ (Supplemental Fig. 4). The mean ± S.E.M phase lag between contacts during co-ripples was 0.07 ± 0.003 rad during waking and -0.12 ± 0.004 rad during NREM.

**Figure 2.**
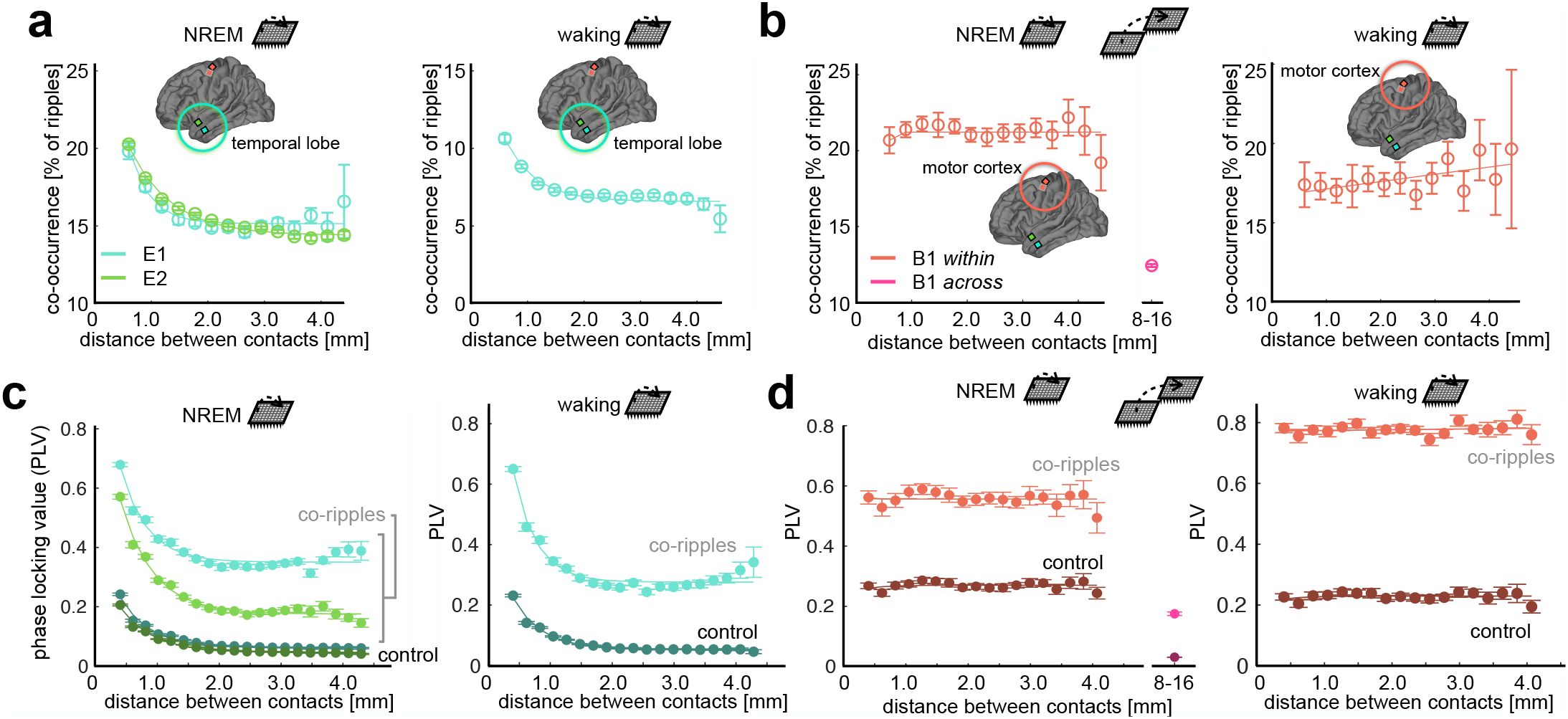
Ripple co-occurrence and phase locking. **a** Mean ± SEM ripple co-occurrence rate as a function of distance in the temporal lobe during NREM (left) and waking (right). The percent of ripples that overlap by a minimum of 25 ms is shown along the abscissa. **b** Same as **a**, but for the motor cortex. Ripple co-occurrence is shown across both Utah arrays in NREM. **c** Co-ripple phase locking value (PLV) over distance during NREM (top) and waking (bottom) in the temporal lobe. PLV for control periods absent of ripples in either channel pair is also shown. Note – PLV does not depend on oscillation amplitude. **d** Same as **c** but in the motor cortex. PLV is also shown across Utah Arrays in NREM.

### Co-rippling increases unit co-firing during waking and NREM

We next tested the prediction of BBS that co-firing across cortical locations is greater when those locations are co-rippling. Indeed, unit co-firing rate during co-ripples that overlapped by at least 25 ms was increased compared to duration matched ripple-absent control periods during NREM (mean ± SEM: 0.655 ± 0.006 Hz vs. 0.185 ± 0.002 Hz, p ≈ 0, one sample two-sided Wilcoxon signed-rank) and during waking (0.909 ± 0.009 Hz vs. 0.479 ± 0.005 Hz, p ≈ 0) for each patient included in this study (Fig 3A,B). Co-firing was increased for unit pairs spanning all distance bins within arrays for NREM and waking. During NREM, the increase in co-firing was maintained for units up to ∼16mm apart across both M1 arrays (0.387 ± 0.009 Hz vs. 0.155 ± 0.004 Hz, p ≈ 0). Within-array co-firing during co-ripple periods increased by 164% for PY↔PY, by 209% for PY↔IN, and by 296% for IN↔IN interactions during waking. In NREM, within-array co-firing increased by 329% for PY↔PY, by 374% for PY↔IN, and by 430% for IN↔IN interactions. Across M1 arrays, co-firing increased by 241% for PY↔PY, by 254% for PY↔IN, and by 254% for IN↔IN interaction (see Supplemental Table 1 for all interaction specific firing rates).

**Figure 3.**
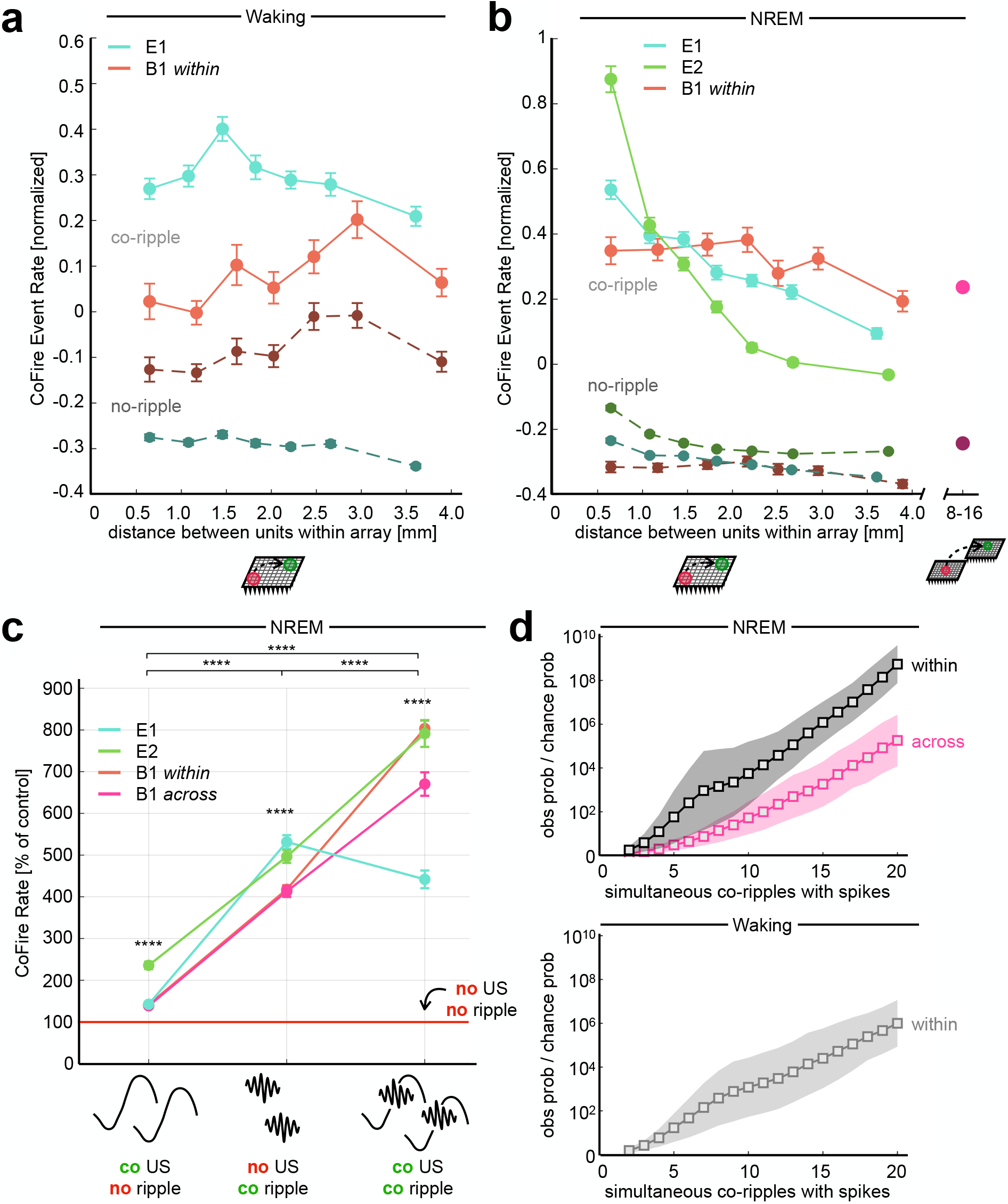
Co-rippling increases co-firing across sub and supra-millimeter scales. **a** Waking co-firing is increased during co-ripples across entire array. Each distance bin contains an equal number of unit pairs. All unit pairs that span across both B1 arrays are grouped into one bin. A co-fire event is defined as a co-ripple period that contains an action potential in both co-rippling sites. Controls are duration matched periods where neither site is rippling. **b** Same as **a**, but for co-firing during NREM. **c** Co-fire rate within 25ms across isolated co-occurring upstates, isolated co-occurring ripples and co-occurring ripples coupled to upstates. Control period is defined as all regions of the recording absent of upstates and ripples (**** p < 0.0001, one sample two-sided Wilcoxon signed-rank test). **d** The observed over chance probability that ripples containing at least one spike co-occur increases with the number of spike containing co-ripples. Top, NREM. Bottom, waking.

### Unit co-firing increases during co-ripples are not dependent on coupling with upstates, but are larger when they do

Given previous reports that ripples tend to couple with slow waves during NREM^10,14^, we quantified how ripple induced co-firing compared with slow-wave induced co-firing, and whether ripple coupling with upstates influences co-firing (Fig 3C). We detected upstates during NREM epochs in all electrodes with at least one detected single unit according to previously described techniques^14^. Upstates had a mean amplitude and density of 178.3 ± 63.1 μV and 10.1 ± 2. min^-1^ across Utah Array contacts. Ripples were defined as ‘coupled’ to an upstate if they occurred within a 100ms window before the upstate peak. Epochs were identified where both ripples and upstates co-occurred on a given electrode pair, where ripples co-occurred without upstates on either electrode, where upstates co-occurred without ripples on either electrode, where neither ripples nor upstates occurred on either electrode (‘baseline’). A co-firing window of 25ms was used for all analyses in this section. Co-ripples without upstates increased co-firing by 375% compared to baseline (mean ± SEM: 1.073 ± 0.023 Hz, vs. 0.226 ± 0.006 Hz, p ≈ 0, one sample one-sided Wilcoxon signed-rank). In contrast, co-occurring upstates without co-ripples only slightly increased co-firing (0.332 ± 0.007 Hz vs. 0.226 ± 0.006 Hz, p ≈ 0). When co-ripples coupled to co-occurring upstates, co-firing was increased an additional 68% compared to isolated co-ripples alone (1.802 ± 0.035 Hz vs 1.073 ± 0.023 Hz, p ≈ 0). Therefore, isolated co-ripples do increase co-firing, and that effect is greater when co-ripples are coupled to co-upstates. The same ripple/upstate co-firing dynamics were observed between units spanning within and across Utah Arrays.

### Co-ripples can increase co-firing beyond that expected from increased overall firing

We next tested if the increase in pairwise co-firing during co-ripples is purely a result of the independent increase in firing rates for each unit during their respective ripples. Specifically, we compared the observed co-firing rate during co-ripples to the co-fire rate during a shuffled co-ripple control (i.e., co-ripple periods where the inter-spike intervals for each unit pair are shuffled 1000 times, α = 0.05). Any co-firing above the shuffled control is therefore more organized than what can be explained by the increase in pairwise firing rates alone. To exclude very sparsely firing cells, we only examined cell pairs that each had a firing rate of 1 Hz during co-ripple periods between their two sites. For all within-array pairs during waking, 6.70% of PY→PY pairs, 5.01% of PY→IN pairs and 5.59% of IN→IN pairs showed increased co-firing within 10 ms (roughly the period of a ripple cycle) compared to co-ripple shuffles. During NREM, the proportion of cell pairs exhibiting highly organized firing increased to 6.7% of PY→PY pairs, 6.5% of PY→IN pairs and 11.2% of IN→IN pairs.

### Co-firing during co-ripples tends to occur in multi-site groups

We previously found with local field potential recordings that the strength of phase-locking between two co-rippling locations strongly increases with the number of additional co-rippling sites. The number of co-rippling sites itself was much greater than expected, and the degree of observed/expected increased exponentially with the number of co-rippling sites^15^. Similarly, here we compared the probability of finding spiking in all N sites if those N sites are all co-rippling, to the expected value if co-firing with co-ripples is independent of the number of sites co-rippling (calculated as the product of the probabilities of each of those sites spiking if it is rippling, regardless of whether other sites are rippling). The observed ratio is ∼1.4x higher than expected if N=2, and ∼10^8^x higher if N=20 (Fig. 3D). Thus, interaction strength increases rapidly with the number of interacting sites.

### Co-rippling increases predictive firing between units up to 16 mm apart

The co-firing analysis reported above only considers the co-occurrence of firing within a 25ms window, without considering the order of spiking, its timing, or a lengthier temporal context. Here, we examine whether the timing of firing by a neuron in one location over an extended period (150ms) better predicts the firing of another spatially separated neuron when their two locations are co-rippling.

To this end, we predicted the firing of each detected unit by every other simultaneously recorded unit based on a coupling filter constructed from their normalized pre-spike pairwise cross-correlograms (CCG). For all prediction analyses, the predicted target neuron is labeled neuron A, while the predicting driver neuron is labeled neuron B. For a given neuron pair, a pre-spike coupling filter between neuron B and neuron A was iteratively constructed, leaving out one action potential in neuron B at a time (mimicking a jackknife bootstrapping technique). The resulting filter was normalized across the pre-spike window such that its sum equaled zero. The removed B spike was fed into the normalized coupling filter, obtaining a B→A prediction measure for that single spike. Any pair of A and B neurons whose average prediction is greater than zero therefore demonstrated consistently organized firing latencies by neuron B within the window prior to spikes by neuron A. Co-rippling vs no-rippling predictive coupling filters were constructed from equal numbers of spikes for each cell-pair to eliminate confounds from increased co-firing during co-ripples (Fig. 4A,B). As with all co-firing quantification, the coupling between neurons detected on the same contact was not examined in this analysis.

**Figure 4.**
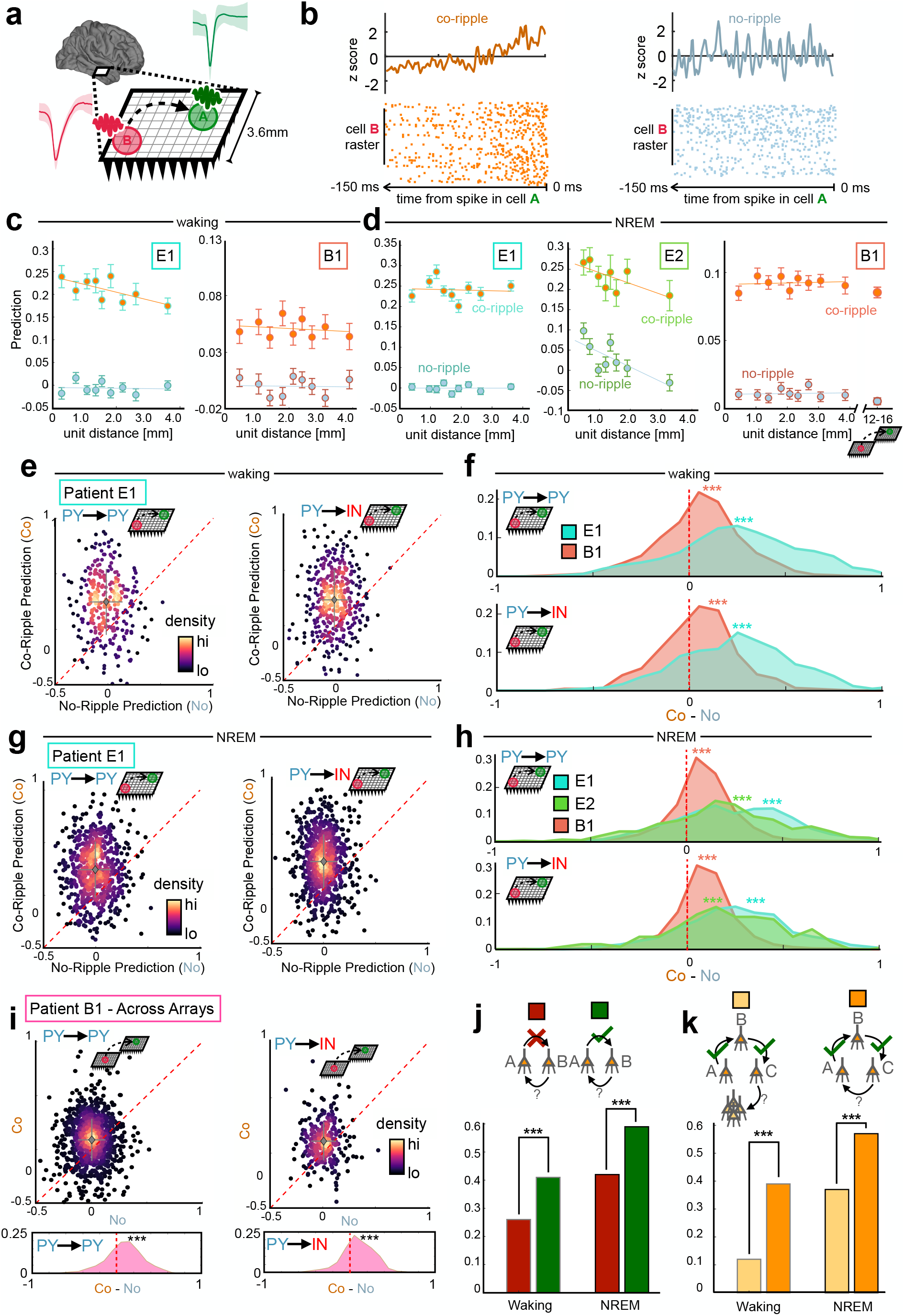
Co-rippling increases pairwise predictive neural firing patterns. **a** Quantification of predictability: The ability of neural activity of a driver neuron in site B to predict firing of a target neuron in site A, typically between when the two sites are co-rippling (‘co-ripple’) versus when neither is (‘no-ripple’). **b** Prediction filter design. Cross correlograms between neuron A and B are computed, smoothed and z-scored within a 150ms window. Relative A-B firing times are fed into the filter, obtaining a prediction value for each action potential in the B neuron. This process is repeated when each site is co rippling and each site is not rippling (controlled for the number of B action potentials). **c** Co-ripple and no-ripple prediction during waking within arrays. Mean ± SEM is shown for distance bins with an equal number of unit pairs. **d** same as **c**, but during NREM and additionally showing prediction between arrays for patient B1. **e** Top, scatter plot of mean within-array co-ripple prediction and no-ripple prediction for each unit pair in patient E1 for PY→PY and PY→IN interactions. Data is shown for waking. Dashed red line is where co-ripple = no-ripple predictability. Bottom, histogram of co-ripple minus no-ripple prediction for each unit pair. **f** Within-array co-ripple minus no-ripple prediction distributions for patients E1 and B1 during waking. **g** Same as **e** but showing prediction during NREM. **h** same as **f** but for all patients during NREM. **i** same as **g** but showing prediction for all unit pairs spanning across arrays in patient B1. **j** Quantifying reciprocal predictive connections connection. Bar graphs show the probability that B predicts A when A does predict B (green) and A does not predict B (red). **k** Quantifying reciprocal connections in ordered triplets. Bar graph shows the probability that cell C predicts cell A (orange) and cell C predicts any other cell in the UA (yellow) in networks when A predicts B and B predicts C. *** p < 1e-4.

In spontaneous waking, we evaluated prediction for 217 PY, 44 IN and 36725 directed cell pairs within Utah Arrays implanted in patients E1 and B1. Of all within-array directed cell-pairs, 4261 (12%) had the minimum of 25 B neuron spikes we required for further prediction analyses. Of the cell pairs that met the B spike inclusion criteria, 68% had greater predictive coupling during co-ripples (p < 1e-10, paired one-sided Student’s t-test on co-ripple vs no-ripple prediction). Furthermore, while prediction strength remained near zero across distance bins in non-rippling periods, during co-ripples it remained significant and above no-ripple levels for unit pairs up to 3.6mm, the longest distance with enough cell pairs (Fig. 4C). Overall co-ripple prediction was higher than no-ripple controls for all unit type interactions and across all patients in waking (Fig 4E,F, Supplemental Fig. 5). Specifically, prediction during co-ripples was stronger in 66% of PY→PY, 70% of PY→IN, 67% of IN→PY and 74% of IN→IN pairs that met the B spike count inclusion criterion (p<1e-10, paired one-sided Student’s t-test for all cell type interactions).

In NREM, we evaluated prediction for 437 PY, 72 IN and 73974 directed cell pairs within Utah Arrays implanted in all patients. Prediction levels during NREM did not substantially differ from waking levels (Fig. 4D). Out of all possible cell pair combinations in NREM, 9476 (13%) met B spike inclusion criteria for further analysis. As in waking, overall co-ripple prediction was higher than no-ripple controls for all unit type interactions (Fig. 4G,H). We found that 76% of PY→PY, 79% of PY→IN, 76% of IN→PY and 82% of IN→IN pairs were higher during co-ripples (p<1e-10, paired one-sided Student’s t-test for all cell type interactions).

In the dual array separated by 7-16 mm, prediction was measured for 120 PY and 27 IN cells during NREM, leading to 4982 total directed cell pairs and 2160 (43%) pairs meeting inclusion criteria that spanned across both arrays. Overall, prediction was comparable across arrays compared to within-array levels in patient B1 (Fig 4D). Prediction was significantly higher during co-ripples (Fig. 4I) in 68% of PY→PY, 72% of PY→IN, 69% of IN→PY and 62% of IN→IN pairs (p = 0.03 for IN→IN pairs, p < 1e-10 for all other cell type interactions, paired one-sided Student’s t-test). Thus, co-ripple facilitation of unit-unit firing prediction is present in both temporal cortex and motor cortex, within and across arrays, as well as in spontaneous waking and NREM. The increased prediction strength during co-rippling vs. no-rippling periods was robust to variations in the parameters used to compute predictive coupling between cells. The difference between co-ripple and no-ripple conditions remained significant across pre-spike coupling filter widths between 50 and 750 ms, and pre-filter baseline window lengths from 0 to 600 ms (Supplementary Fig. 6).

### The degree of unit phase locking to underlying ripple modulates predictability

A key tenet of the BBS hypothesis is that the phase of synchronized gamma oscillations modulates neural excitability in different cortical locations at precise timescales, thus facilitating their interactions. We therefore sought to examine whether the relationship between neural firing and LFP oscillation phase during co-ripples influences pairwise predictive coupling. We first quantified the degree to which the level of phase locking between neural firing and ripple phase modulated prediction for pairs of neurons that are significantly coupled during co-ripples (Fig 5. A,B). In NREM, the phase locking value (PLV) between the driver neuron (neuron B) and ripple phase was positively correlated with prediction level during co-ripples (r_B_ = 0.38, p < 1e-10, Pearson’s correlation). The same relationship was true between the PLV between the target neuron (neuron A) and ripple phase (r_A_ = 0.31, p < 1e-10). The strength of correlation was slightly greater than the sum of its parts when the PLV from neuron A and B were summed (r_A+B_ = 0.44, p<1e-10), suggesting a possible synergistic effect of phase-locking in both the driving and target regions. Notably, the subset of neuron pairs that spanned across both arrays in patient B1 exhibited the same PLV-prediction relationships (r_A_ = 0.22, p < 1e-5, r_B_ = 0.29, p < 1e-5, r_A+B_ = 0.34, p < 1e-5).

**Figure 5.**
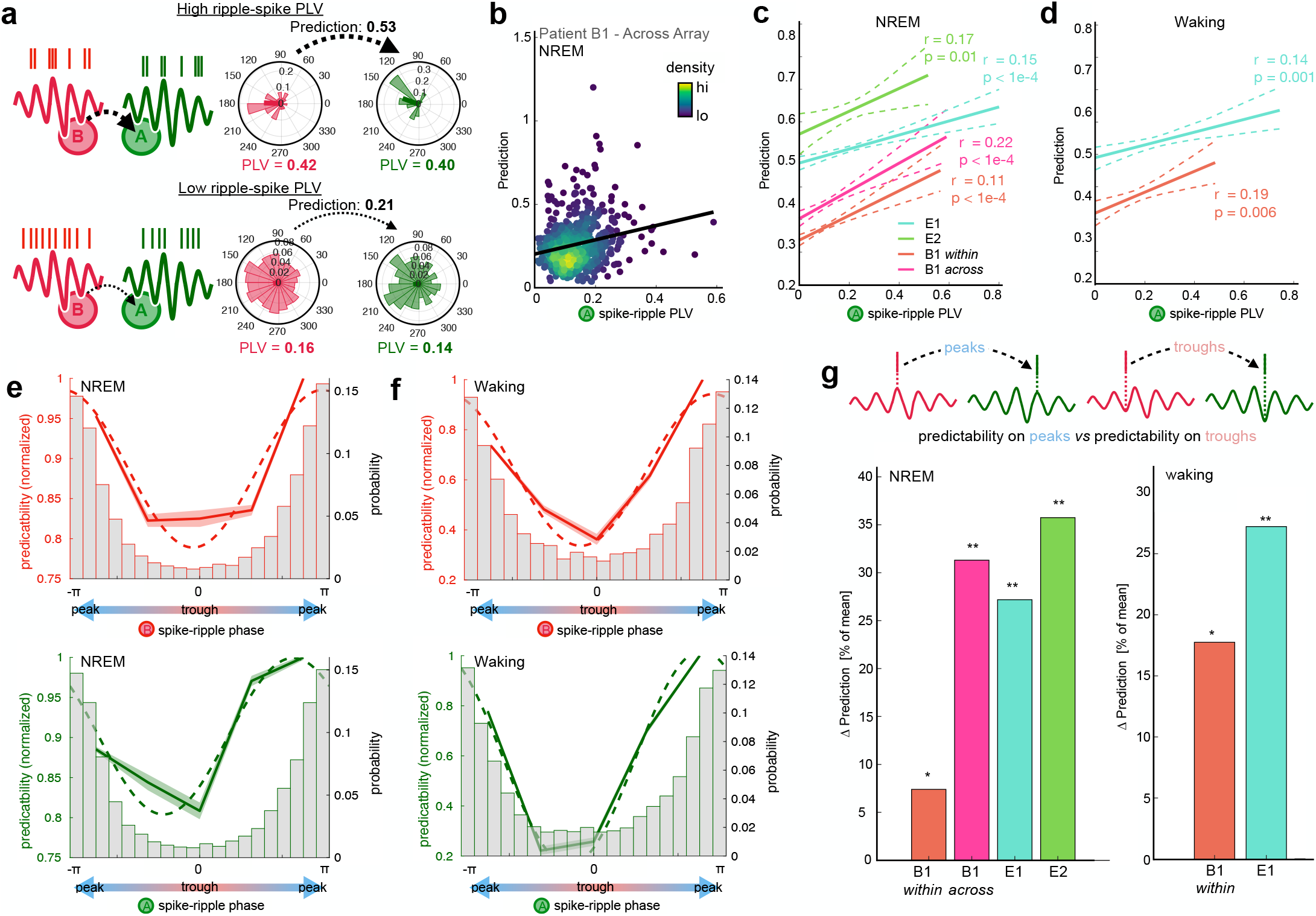
Spike-phase relationships modulate prediction during co-ripples. **a** Schematic depicting the prediction for a high PLV (top) and a low PLV (bottom) neuron in the driving (i.e., B) and target (i.e., A) locations. **b** Co-ripple prediction is positively correlated with the level of phase locking between the A neuron and the ripple. Each dot represents a unit pair that spans across arrays in patient B1 (N = 560 pairs, r = 0.29, p < 0.01). **c** Prediction relationship with A spike-ripple PLV in all subjects. Dashed line shows 95% confidence interval of bootstrapped distribution (N = 1000 iterations, with replacement). **d** Same as in **c** but during waking. **e** Top, prediction as a function of preferred B spike-ripple phase across all subjects. Bottom, prediction as a function of preferred A spike-ripple phase across all subjects. Normalized mean prediction is shown across all unit pairs for 5 equally spaced bins from -π to π. Dashed line shows best sinusoidal curve fit. Histogram shows the distribution of ripple phase preference for B (top) and A (bottom) neurons. **f** Same as **e** but during waking. **g** Change in prediction when both A and B spikes are on ripple peaks vs when both are in ripple troughs. For each subject, Δ prediction is computed for all A-B pairs that are positively predictive in co ripples. Left, NREM. Right, waking. (*p < 0.05, ** p <0.01, one-sided paired Student’s t-test). The Spearman rho values are displayed for each individual patient. Some traces do not reach 100% since the periods of the recording that contained exactly that number of co-rippling channels were below the minimum duration threshold for analysis (1 second). ** p < 0.01, *** p < 0.001 one-sided one-sample Student’s t-test.

During waking, ripple-phase PLV for both the driving (r_B_ = 0.42, p < 1e-10) and target (r_A_ = 0.36, p < 1e-10) neural activity were also positively correlated with prediction. The correlation was also slightly stronger during waking when the driver and target neuron PLVs were summed (r_A+B_ = 0.50, p < 1e-10). The PLV relationships with prediction were significant for each individual patient during waking and NREM (Fig.5 C,D). This suggests that the consistency between neural activity and oscillation phase during co-ripples influences the magnitude of their coupling.

### Prediction is increased for spikes occurring on ripple peaks

To further test the prediction of the BBS hypothesis that phase modulation promotes neural interactions, we examined whether the specific phase preference (i.e., peak or trough) of the driver and target neurons influences pairwise prediction. Across all cell pairs, we measured the circular mean ripple phase preference and average prediction for both the driver and target neurons during co-ripples. Prediction was significantly sinusoidally modulated by ripple phase in both NREM and waking (sinusoidal fit significance determined from 1000 shuffled permutations of prediction – phase pairings, Supplemental Fig. 7). In both behavioral states and for both driver and predicting neurons, phase preferences around the ripple peak showed higher prediction compared to phase preferences around the ripple trough (Fig. 5E,F). The same modulation of prediction by phase was observed for unit pairs within and spanning across arrays. This indicated that neuron pairs with higher predictive coupling tended to have both driver and target neurons with phase preferences at the peak of the ripple.

To gain further insight into the influence of ripple phase on prediction, we next analyzed the subgroup of neuron pairs that had positive prediction during co-ripples. Within each of the co-ripple modulated cell pairs, we compared the mean prediction in the subset of cases when both driver and target spikes occurred on their respective ripple peaks (double peaks) to the subset of cases when they occurred on their respective ripple troughs (double troughs). Within each co-ripple modulated cell pair, prediction increased by 21% for double peaks compared to double troughs during NREM (p < 1e-10, paired one-sided Student’s t-test, N_pairs_ = 7535), and 25% during waking (p = 2e-4, paired one-sided Student’s t-test, N_pairs_ = 2014). Double peaks had greater prediction compared to double troughs for all individual subjects, within and across arrays (Fig. 5G). These results suggest that the ripple peak reflects a more favorable environment for the interaction of cells spanning up to 16 mm apart.

### Co-prediction is enriched for reciprocal connections

The above analyses of prediction of firing by one cell by another during co-ripples in their sites were all focused on directed pairs of neurons. We also tested if neurons were mutually predictive. This direct return, where both A→B and B→A significantly predict each other’s firing during co-ripples represents second order recurrent activation (Fig. 4J). In NREM, for all significantly predictive A→B cell pairs, 59% of the reciprocal B→A connections were predictive, compared to 42% when A→B was not significantly predictive (p < 1e-10, X^2^ = 119.0, df = 1). In waking, for all predictive A→B cell pairs, 41% of the reciprocal B→A connections were predictive, compared to 26% of A→B pairs that were not significantly predictive (p = 5e-10, X^2^ = 38.7, df = 1).

As noted above (Fig. 3D), co-rippling typically engaged more than 2 sites. We thus explored whether recursive co-prediction networks extended beyond two cells by examining prediction across ordered triplets of neurons (Fig. 4K). Given a network where A→B and B→C are significantly predictive, we asked what the probability was that C significantly predicted the firing of A (C→A) vs any other cell in the array (C→XX). In NREM, the recursive C→A connection was predictive in 57% of cases, while 37% of C→XX connections were predictive (p ∼ 0, X^2^ = 19912, df = 1). In waking, the C→A connection was predictive in 39% of cases, while the C→XX was predictive in 12% of cases (p ∼ 0, X^2^ = 7402.3, df = 1). Thus, it appears that ordered sets of neurons where the activity in one predicts the other during co-rippling, concatenate and often form recurrent networks of prediction, at least on the scale of a UA.

### Co-rippling modulates the expression of neuronal assemblies (Fig. 6)

Neural coding is unlikely to be fully explained by pairwise interactions alone. Recent work in humans^28^ suggested that groups of neurons that have significant co-firing tendencies can be categorized into cell assemblies whose expression is linked to episodic memory performance. We next evaluated whether integrated neural assemblies were detectable in our data and whether co-rippling across multiple electrodes facilitates their expression. We detected cell assemblies during NREM and waking according to previously described methods^28,29^. We divided neural spike trains from NREM and waking into 100 ms bins (roughly the duration of a ripple event) and normalized the binned firing rates for each neuron across the recording session. Using a combination of principal component analysis (PCA) followed by independent component analysis (ICA), we identified groups of neurons with significant coactivation patterns. This produced an array of component weights across all neurons for each assembly, indicating how strongly a neuron contributed to the expression of a given assembly. Highly contributing cells were identified as key member neurons of the assembly if their ICA weight exceeded 1.5 standard deviations above the mean weight (Fig. 6A). The expression of each assembly can be determined by projecting the binned z-scored firing matrix onto the assembly weight vector. We defined assembly activation events when expression exceeded the 95th percentile across the recording (Fig 6B).

**Figure 6.**
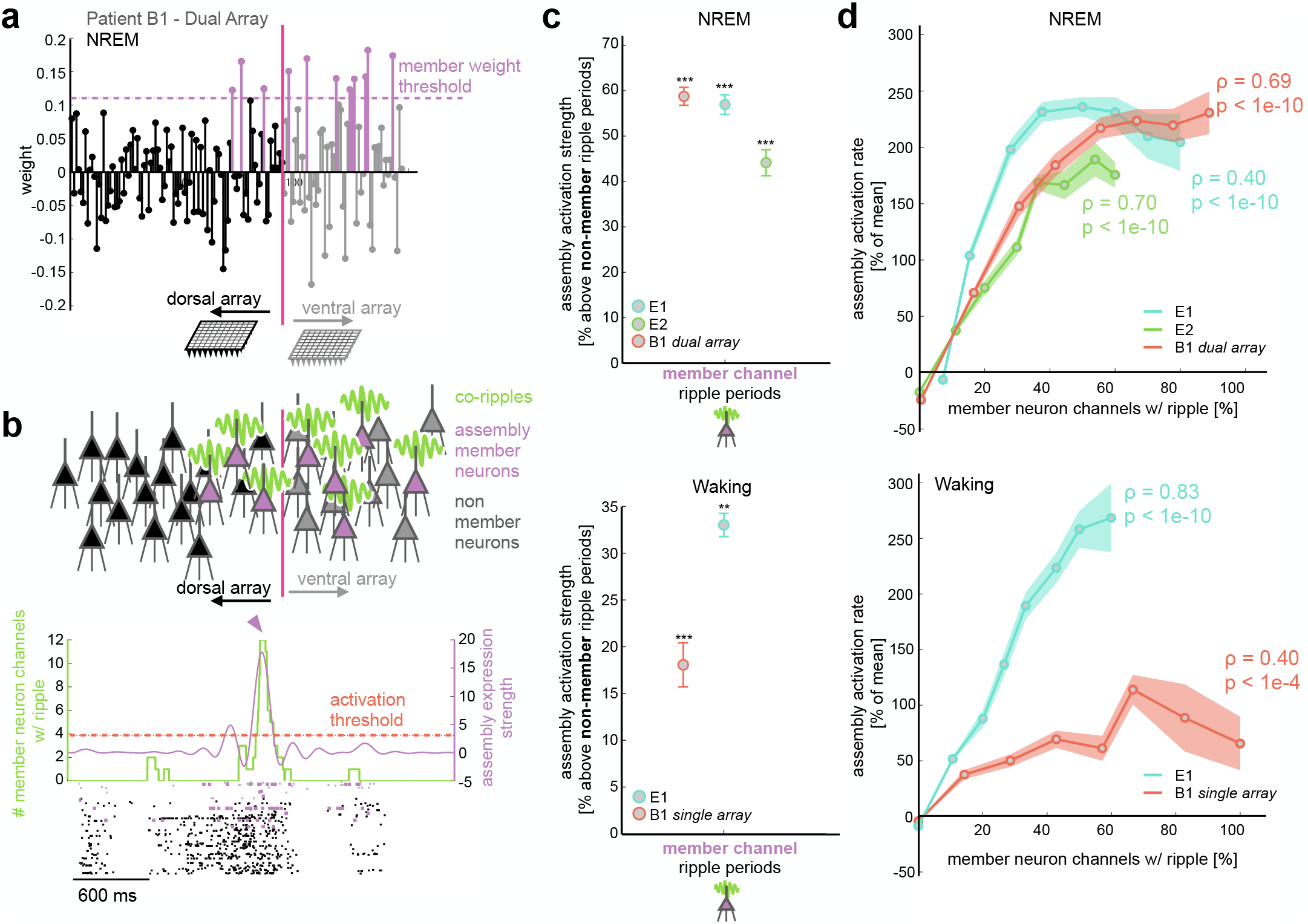
Co-ripples facilitate expression of neuronal assemblies. **a** Example assembly with identified member neurons that span across both arrays in subject B1 during NREM. **b** Top, schematic depicting assembly member neurons in sites that are co-rippling. Bottom, example of activation of the assembly in **a** when all member neuron sites have a detected ripple. Assembly strength (blue curve) and ripple count (green curve) are superimposed on the spike raster of both arrays in subject B1. Colored spike raster depicts cells that were labeled as assembly members. Horizontal dashed red line displays the 95%ile activation threshold, and the blue carrot marks the time where the displayed assembly is activated. **c** Mean ± SEM assembly strength above control during periods where member neuron channels have a detected ripple. The control is defined as periods when non-member neurons have a detected ripple. NREM is shown on the top and waking on the bottom. **d** Mean (circle marks) ± SEM (shaded region) assembly activation rate as a function of the proportion of member neuron electrodes that contain a detected ripple. Top, results in NREM for all patients (spearman rho = 0.59, p < 1e-10). Bottom, results in waking using data from the medial array in patient B1 and the array in E1 (Spearman rho = 0.50, p < 1e-10).

We identified 76 total assemblies during NREM and 40 total assemblies during waking (23 ± 4 per patient per behavioral state). Each assembly had a mean ± std. dev. of 7.9 ± 3.2 member neurons during waking and 10.6 ± 2.1 member neurons during NREM. Of note, in patient B1 during NREM (when data was available for both Utah Arrays), key member neurons often spanned across dorsal and ventral arrays (Fig. 6A,B, Supplementary Fig. 9). We compared the mean expression strength during periods when at least one member neuron channel was rippling to periods when no member neuron channels had a detected ripple, but at least one non-member neuron channel did. Mean assembly strength was higher during member rippling periods compared to non-member ripple controls (Fig. 6C) during waking (24% increase, p = 1e-4, one-sided paired Student’s t-test) and NREM (59% increase, p = 5e-6, one-sided paired Student’s t-test). This suggests that co-ripple networks are associated with the precise selection of co-active cells. We next measured the activation rate of each assembly as a function of the percent of member neuron channels that were co-rippling. Assembly activation rate was positively and monotonically correlated with the proportion of member neuron channels that were engaged in a co-ripple in NREM (p = 1e-10, Spearman rho = 0.57) and in waking (p < 1e-10, Spearman rho = 0.52) for all patients (Fig 6D).

## DISCUSSION

The mechanisms by which the cortex binds the activity of neurons separated by relatively large distances remain largely unknown. Bursts of high frequency oscillations (‘ripples’), have recently been shown to possess some of the characteristics necessary for a role in cortical neural binding^14,15,23^. They are ubiquitous and consistent in frequency and duration across cortical areas, and across waking and sleep. They have a strong tendency to co-occur and phase-lock even at long distances, and phase-modulate firing of local pyramidal and interneurons. While these properties are prerequisites for Binding-By-Synchrony, they do not establish that the basic function of binding, integration of cortical activity, is facilitated by co-rippling. Such was the goal of the current study.

Local field potentials and single units were recorded during NREM sleep and waking with four 96-channel microarrays, each implanted in a 16mm^2^ patch of human supragranular cortex. Single arrays were implanted in the lateral temporal neocortex of two epileptic patients, and two arrays separated by ∼12 mm were implanted in the precentral gyrus of a tetraplegic patient. Across patients, cortical areas, and states, we found increased integration of neuronal firing in different cortical locations when those locations co-rippled, using three measures: co-firing, co-prediction, and assembly expression. These measures of the interaction between the firing of cells in different locations are sensitive to increasing complexity and duration of the predicting spike train. Co-firing only requires that cells fire within 25ms of each other; co-prediction is sensitive to the temporal structure of firing increases and decreases by the predicting cell in the 150ms prior to the predicted cell’s action-potential; assembly expression detects participation in an extended network involving multiple cells. The similar results obtained with all three measures across areas and states suggests that increased interaction of cortical neuronal firing during co-ripples is a robust phenomenon.

The central mechanism underlying Binding-By-Synchrony is the synchronous modulation of cell excitability in different locations, whereby action-potentials triggered by depolarization in one location would be effective at triggering action-potentials in another because of its depolarized state^2^. In this view, ripple phase indicates the membrane potential of pyramidal cells, and ripple synchronization would thus indicate the synchronization of excitability of cells in the co-rippling locations. This would result in co-firing, and more generally in the selection of neural networks that engage the particular locations that are co-rippling. Accordingly, we found that co-ripples frequently occur within and between Utah arrays, where neurons often fired in phase with the local ripple^15^. Critically, we found that the level of this phase-locking is correlated with the ability of neuronal firing in one location to predict the firing in another location. Specifically, prediction is enhanced when both predicting and predicted spikes occur on the ripple peak, as posited by Binding-By-Synchrony.

Some have suggested that ripples do not actively facilitate transcortical integration but are simply a byproduct of nonspecific activation^4,5^. However, co-rippling still enhanced co-firing when compared to co-firing to the same spikes with their inter-spike intervals shuffled, indicating that the precise relative timing of the spikes in the two locations is critical. Similarly, our method of cross-location prediction balanced the firing in no-rippling and co-rippling predictors, but still found increased prediction during co-rippling. These controls, and the strong effects of phase on prediction, demonstrate that the effects of co-ripples extend beyond those due to co-activation.

The arrival of an action-potential from one location to another not only has a higher probability of evoking a spike in the post-synaptic cell if it arrives on the depolarized ripple phase, it also augments that depolarization, and thus contributes to the ripple. Furthermore, we found that co-ripple-enhanced prediction is reciprocal (if neuron A predicts B then B likely predicts A) and recurrent (if A predicts B which predicts C, then C likely predicts A), Thus, co-ripples could initiate positive feedback loops that sustain and possibly augment specific patterns. These considerations suggest that a strongly positive relationship may be found between increasing levels of co-rippling and measures of firing interaction. Indeed, we found a large increase in co-firing during co-ripples with increasing co-rippling and co-firing in other sites. Similarly, the level of assembly activation increases steeply with an increasing proportion of sites that are co-rippling. Thus, increased firing and phase modulation may be better viewed as synergistic and mutually reinforcing rather than as alternatives.

Most previous studies of ripples were of hippocampal ripples in rodents during NREM sleep where they play a critical role in facilitating cortical consolidation, in conjunction with slower waves including cortical upstates^7,8^. We previously found that cortical ripples in humans are strongly related to upstates during NREM^14^. Here we found that the co-occurrence of either upstates or ripples alone increases co-firing, with the increase due to co-ripples about twice that to co-upstates. However, when both co-upstates and co-ripples occur together, the increase in co-firing is much greater than either alone. The window for these increases is 25ms, the window when Spike Timing Dependent Plasticity modulates synaptic strength^30^. Thus, these results reinforce the potential role of nested oscillations in promoting consolidation during sleep.

There are several limitations to this study. First, data is provided from only three patients, two of whom have long-standing epilepsy. However, no interictal spikes or seizures were observed in the epochs studied here, and no pathology was visible in this location in neuroimaging. Critically, ripple characteristics, co-occurrence and modulation of neural activity were similar between these two patients with epilepsy and the third patient who is paralyzed but has a healthy motor cortex. Since this appears to be the first work quantifying ripples with intracranial recordings in healthy human cortex, it provides strong evidence that ripples are not epileptic phenomena.

The second limitation is anatomical: we only recorded from supragranular layers of limited regions in two cortical areas. Multi-patch recordings in humans^31^ and anatomical studies in primates^32^ suggest that horizontal connections may be more intense in layers 2/3, raising the possibility that integrative processes may be different in other layers. Each Utah array subtends ∼16mm^2^ of cortical surface, ∼0.007% of the total, and even in the patient with dual arrays the longest distance separating neurons is ∼16mm, ∼7% of the longest cortico-cortical streamline distance estimated from DTI^33^. Thus, it is important to confirm our results in larger and more distributed datasets. However, we would note that the effects we describe decrease minimally or not at all over the distances we sampled, and in previous work the probability of co-rippling and level of phase-locking was largely maintained for over 200mm^15^.

The third limitation is the lack of a behavioral task. While binding of course occurs during spontaneous mentation, and ripples have important putative functions during sleep, this limitation, together with the lack of recordings from early visual cortex (for clinical reasons), as well as species differences, renders it difficult for us to compare our results directly to most experimental literature testing Binding-by-Synchrony, which are typically in visual tasks with recordings from early visual cortices in cats or rodents.

In summary, we provide evidence that co-rippling in different cortical locations enhances the integration of their constituent neuronal firing. This finding was replicated with co-firing, co-prediction, and neural assembly activation, in three subjects, two cortical areas, and sleep and waking, for putative pyramidal and interneurons. When two cortical locations co-ripple, we find a corresponding increase in pairwise neural co-firing and prediction for units detected in those two locations. Co-ripple facilitated prediction is strongly modulated by ripple phase, persists at the longest separations sampled (16mm), and remains when firing rate increases during ripples are controlled for. These results support in humans key predictions of the Binding-by-Synchrony model: that ripples co-occurring in different cortical locations facilitate the integration of their neuronal firing through a mechanism involving phase modulation.

## Methods

### Participants and data collection

The data used in this study was acquired in two types of patients. The first category consisted of adult patients (Table 1) with focal, pharmaco-resistant epilepsy undergoing 4–21 days of continuous invasive EEG recordings for the localization of seizure foci prior to resection. While undergoing clinical recording these patients were also implanted with a Utah Array microelectrode recording within the region of therapeutic resection (Fig.1a; Utah Array–© 2020 Blackrock Microsystems, LLC). The Utah Arrays were implanted in accordance with the review of Institutional Review Board (IRB) of Partners HealthCare, the parent institution of Massachusetts General Hospital and Brigham and Women’s Hospital. In all cases, the UA implant location was resected to gain surgical access to the seizure focus in deeper structures. The resected tissue in which the Utah Array was implanted was determined not to be an epileptogenic in any of the patients analyzed in this study. Furthermore, no seizures occurred during the periods analyzed in this study.

The second category included a patient with a cervical spinal cord injury participating in the BrainGate clinical trial (www.ClinicalTrials.gov; Identifier: NCT00912041). The patient had a history of traumatic brain injury and encephalomalacia in the left inferior frontal lobe, but otherwise had a healthy cortex. The Utah Arrays were implanted in the patient with permission granted by the U.S. Food and Drug Administration (Investigational Device Exemption #G090003) and the IRBs of Massachusetts General Hospital, Providence VA Medical Center, and Brown University. The patient, (identified as B1 in this study and T11 in others^20^, gave informed consent to the study and publications resulting from the research. We analyzed neural recordings from the second set of two separate multiday sessions. Details of the recording setup are described in previous work^20,34-37^. Array placement was performed 427 d before the recording session analyzed in this study.

### Electrodes and localization

The Utah Array is a 10 × 10 microelectrode grid, with corners omitted, that has 400μm contact pitch (Fig.1a). Each silicon probe is 1 or 1.5 mm long and 35– 75μm wide at its base, tapering to 3–5μm at the tip, and is insulated except for the tip, which is platinum-coated. Each epilepsy patient in the study had one array implanted into the superior or middle temporal gyrus, in a region that was outside of the epileptogenic focus, but which had to be removed to gain surgical access to the focus. As part of the ongoing BrainGate clinical trial, the patient with tetraplegia had two arrays implanted in the left precentral gyrus. The two probes were separated by a center-to-center distance of 12 mm (contact distance range: 7 – 16 mm). All probes were placed under direct visualization perpendicular to the cortical surface.

Data were acquired at 30 kHz sampling (Blackrock Microsystems), from 0.3 to 7.5 kHz. Data were subsequently low-passed at 500 Hz and down-sampled to 1 kHz for the LFPs. Data were saved for offline analysis in MATLAB 2022b (MathWorks). Channels were excluded when there were large amounts of noise, or no units detected. Out of the 96 recording channels the mean number excluded from analysis was 35.25 (range: 11–66). The 1 kHz data was distance-referenced to a distant subdural contact.

### Sleep staging and data selection

Epochs included in the study did not fall within at least 1 h of a seizure and were not contaminated with frequent interictal spikes (IISs) or artifacts. NREM periods were selected from continuous overnight recordings where the *δ* (0.5-2 Hz) analytic amplitude from the cortical channels was persistently increased. Sleep epochs were confirmed by visual inspection to have normal appearing downstates, upstates, and spindles. Waking periods were selected from continuous daytime recordings that had persistently low cortical *δ* as well as high cortical α (8-12 Hz), β (20-40 Hz), and high γ (70-190 Hz) analytic amplitudes.

### Ripple Detection

Ripple detection was performed on the data on each Utah Array contact down sampled to 1000Hz in the same way separately for all states, based on a previously described methods^14,15^. Data were band-passed with a Butterworth filter at the rippleband (70-100 Hz, forward and reverse for zero-phase shift, sixth order), z-scored across wake and NREM epochs separately, and an initial set of candidate events were identified as containing at least three rippleband cycles with a z-score > 1. Candidate events were further sub-selected if the maximum z-score of the analytic amplitude of the rippleband was greater than 3. Adjacent ripples within 25 ms were merged. Ripple centers were determined as the maximum positive peak in the rippleband bandpass. To compute ripple onsets and offsets, the z-scored rippleband analytic amplitude was first smoothed using a sliding gaussian kernel of 100 ms. Onsets and offsets were marked when the smoothed rippleband amplitude envelope fell below a z-score of 0.75. In patients with a history of epilepsy, ripples were excluded if the absolute value of the 100-Hz high-pass z-score exceeded 7 or they occurred within 2 seconds of a ≥ 3 mV/ms LFP change. Ripples were also excluded if they fell within ±500 ms of putative interictal spikes, detected as described in Dickey et al.^14^ Events that had a single prominent cycle or were also excluded if the largest valley-to-peak amplitude in the broadband LFP was 2.5 times greater than the third-largest. For each channel, the mean ripple-locked LFP was visually examined to confirm that there were multiple prominent cycles at ripple frequency, and the mean time-frequency plot was examined to confirm there was a distinct increase in power within the rippleband. In addition, multiple individual and co-occurring ripples in the broadband LFP and rippleband bandpass from each channel were visually examined to confirm that there were multiple cycles at ripple frequency without contamination by artifacts, unit spiking bleed or epileptiform activity.

### Spike detection and sorting

Spike detection and sorting was accomplished using the automated WaveClus 3.0 MATLAB software package^38^. The 30 kHz data recorded from each electrode contact was bandpassed at 300–3000 Hz with an 8th order elliptic filter with a pass-band ripple of 0.1 dB and a stop-band attenuation of 40 dB. Putative unit spikes were detected when the filtered signal exceeded 5 times the estimated standard deviation of the background noise^39^. The spike threshold was therefore computed as:

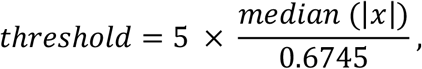

where, x is the 300–3000 Hz filtered data. For each detected spike, 64 samples were saved (20 samples before and 44 samples after the peak of the spike waveform). Features of the resulting waveforms for each spike are decomposed into 64 coefficients using a Haar wavelet. The resulting features are grouped using superparamagnetic clustering (SPC), an unsupervised approach which relies on nearest-neighbor interactions^40^. All clusters detected by the automated approach were manually curated, merging clusters that had similar waveform shapes/amplitudes, and splitting clusters that were clearly comprised of two separate units.

### Single unit classification, quality, and isolation

We used the CellExplorer^41^ waveform classification tool to manually characterize units into putative pyramidal (PY), putative interneuron (IN) and multi-units. Clusters that contained a significant proportion of inter-spike intervals (ISI) within the 3 ms refractory period and/or that had a peak signal-to-noise ratio (SNR) less than 2 were classified as multiunits^42,43^ and excluded from all further analysis in this study. Supplementary Fig.1AB shows the ISI violation and SNR distribution of classified single units. There were cases where multiple single units were classified on a single electrode contact. To determine the degree to which units on the same channel were separated, we measured the pairwise projection distance in units of standard deviations^44^ (see Supplementary Fig.1c for projection distance distribution).

PYs fire at lower rates, frequently burst, and have a broad waveform recovery phase, whereas INs typically fire at higher rates, with infrequent bursting, and have quick waveform recovery. We classified each single unit based on these general characteristics that have been used to separate unit types in rodents^45^ and humans^46^. We estimated waveform recovery using valley-to-peak time and the bursting index was defined from the auto-correlogram using a method described in Royer et al.^47^. PYs were identified as single units with large peak-to-valley times and high bursting indices, while INs had low peak-to-valley times and low bursting (Fig. 1c). While the extracellular waveform only provides indirect insight into neuronal subtype, they have been supported in previous studies to distinguish PYs from INs^48^. Unit stability in multi-day Utah Array recordings has been shown to be robust for units detected 4 days after implantation^49^.

### Removal of spike influence from the LFP

Although extracellular waveforms occur on much faster timescales than oscillatory activity analyzed in the LFP, spikes are large amplitude events that have been shown to often bleed into the LFP and contaminate spike-LFP relationships, especially at higher LFP frequencies^50^. To mitigate this contamination, we subtracted out the mean waveform of each detected unit at all spike times in the raw 30kHz data, before down sampling and filtering the LFP. This template subtraction method successfully removed the direct influence of unit activity on the broadband 1kHz LFP and the filtered rippleband (Supplementary Figure 3).

### Upstate detection and analysis

To evaluate unit spiking during ripples that occurred in association with upstates, we detected upstates and downstates using a previously described method^51,52^. The LFP from each channel was bandpassed at 0.1–4 Hz and consecutive zero crossings within 0.25–3s were detected. The top and bottom 20% of amplitude peaks between zero crossings were then selected. The average high gamma (70–190 Hz) analytic amplitude was found within ±100 ms of each peak. Since the polarity of downstates vs. upstates for each channel was not known a priori, we determined whether the average peak-locked high gamma envelope was higher for positive vs. negative peaks foreach channel and assigned the polarity of upstates (more high gamma) vs. downstates (less high gamma) accordingly. Ripples that coupled with upstates were identified as those where the ripple center occurred within 150 ms of the upstate peak^14^. All upstate-ripple co-fire analysis defined co-firing as spikes from two cells not detected on the same electrode occurring within 25 ms of each other, the window for spike timing dependent plasticity^30^.

### Unit co-firing during co-ripples

Co-ripple periods were identified when there were co-occurring ripples on two channels that overlapped by at least 25 ms. Only the region of ripple overlap was considered as part of the co-ripple period. A co-fire event was defined as a co-ripple period where a spike is measure in both channels. Non-ripple periods were duration matched and selected from regions of the recording where no ripples were present in either channel, with 1 second padded before the onset and offset of every ripple event. Co-firing during co-ripples within 10 ms was also compared to a shuffled control, where the ISIs of each unit were shuffled across all concatenated co-ripple periods 1000 times. Any unit pair with co-firing above the 95^th^ percentile of the shuffled control was considered significant.

### Calculation of pairwise predictive coupling

The statistical predictive coupling between cells was measured using the cross-correlograms (CCGs) between two units. We defined the target neuron as neuron A and characterized the relative spike history of a driving neuron B during co-ripple periods. For each action potential in A that occurred during co-ripples, we measured the spiking in B during a 150 ms pre-spike window in trials where at least one B spike occurred during a ripple on channel B. We next implemented a jackknife approach and constructed the B-A CCGs leaving out one B spike a time. Each CCG was gaussian smoothed using a 5 ms temporal window, which was chosen to be narrow enough not to obstruct the precise relationship between the CCG and the ripple-band phase. Finally, the coupling filter was constructed by z-scoring the smoothed CCG to ensure it summed to zero and positive values reflected substantially organized firing latencies within the filter window. The relative time of the removed B spike was plugged into the filter, obtaining a value for how well that spike predicted a future spike in A. The mean filter output for all B spikes was averaged to compute an overall prediction value between B and A.

Prediction during co-ripples was compared to no-ripple periods where neither A nor B had a detected ripple. The control filters were constructed from a random subset of no-ripple B-A spike history trials that contained the same number of B spikes as the co-ripple condition. Since the number of trials in the control often exceed the co-ripple condition, the co-ripple B spike firing rate was usually higher than controls. To examine the effect that increased firing rate had on prediction, we lowered the co-ripple firing rate below no-ripple levels by removing trials from the co-ripple condition until the firing rate was comparable to the control. Decreasing co-ripple firing rate did not substantially impact the magnitude of co-ripple prediction (Supplementary Figure 8).

### Parameter sweep of prediction method

To validate our choice of parameters, we performed a sweep of the temporal window parameters used in the prediction analysis. We varied the width of the filter from 50 to 700 ms and found that co-ripple prediction remained significantly greater than no-ripple controls (paired one-sided Student’s t-test, α = 0.001) for all subjects in all behavioral states for filter widths > 50 ms (Supplemental Figure 5). We also z-scored across an extended pre-filter baseline window (range: 0 – 600 ms, see Supplemental Figure 5), resulting in coupling filters that did not sum to zero and reflected the increase in co-firing during the 150 ms filter window compared to the baseline period. This approach amplified the substantial increase in co-ripple mediated co-firing between 0 and 150 ms, but partially obfuscated more precise temporal tuning between B and A during co-ripples.

### Assembly detection

We identified neural assemblies using previously described approach applied to human^28^ and rodent^29^ data to detect groups of co-active neurons from spiking data. We first divided spikes into 100 ms bins and z-scored the resulting firing rate matrix to account for neurons with different baseline rates. The bin size of 100 ms was chosen as it roughly represents the duration of a ripple event (Supplementary Fig. 2). We detected co-active assemblies using principal component analysis (PCA) of the z-scored firing rate matrix, and selected significant assemblies from principal components that contained eigenvalues above the bounds defined by the Marchenko–Pastur law^55^.

To minimize the mixing of true cell assemblies, we unmixed the PCA components using independent component analysis (ICA). We first projected the z-scored firing matrix onto the significant assembly principal components that exceed the Marchenko–Pastur bounds:

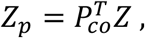

Where *Z* is the z-scored firing matrix, *P*_*co*_^T^ is the transpose of the significant principal components and *Z*_*p*_ is the projection of the firing matrix onto PC space. We use the MATLAB fastica script to solve for the separating matrix *U* of *Z*_*p*_. The separating matrix *U* is then used to rotate the significant PCs to find the optimal assembly patterns:

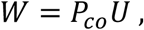

where W contains the final weights for each neuron and for each significant assembly. The magnitude of the weight indicates the degree to which each neuron contributes to a given assembly. We defined significant member neurons for each assembly as neurons whose weight exceeds 1.5 standard deviations above the mean for that assembly.

The time resolved strength of each assembly can be determined using the z-scored firing matrix and the neuron weights:

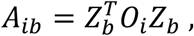

where *A*_*ib*_ is the strength of assembly *i* within time bin *b, Z*_*b*_ is the z-scored firing rate for all neurons in time bin *b* and *O*_*i*_ is the outer product of the neuron weight vector for assembly *i*. We defined assembly activation as time bins where the activation strength exceeded the 95^th^ percentile across the recording. As expected, member neuron firing was significantly increased during assembly activation periods compared to non-member neurons (See Supplementary Fig. 9).

### Statistical analyses

To determine the number of PYs and INs that had significant phase modulation within the 70-100 Hz band, a binomal test was performed between the number of spikes from each unit occurring within 0±π/2 and π±π/2 radians (expected value = 0.5, α = 0.05). To evaluate ripple co-occurrence and PLV vs. non-ripple periods, we used a paired two-sided t-test.

Unit spike rates during ripples vs. non-ripple epochs were compared by using a one sample two-sided Wilcoxon signed-rank test (a non-parametric test was used due to the non-gaussian distribution of firing rate data). Likewise, unit co-firing rate during ripples was compared to duration and number matched non-ripple events using a one sample two-sided Wilcoxon signed-rank test. Co-firing rate within a 25 ms window was also compared between ripples coupled to upstates, isolated ripples, isolated upstates and epochs free of ripples and upstates using a one sample two-sided Wilcoxon signed-rank test. To compare pairwise co-firing during co-ripples vs. shuffled co-ripples, the ISIs of the spikes of both units during all co-ripple periods were randomly shuffled 1000 times. The p-value was calculated as the percent of co-firing within 10 ms that occurred above the 1000 randomly shuffled times.

The significance of co-ripple prediction was determined using a paired one-sided Student’s t-test of the mean co-ripple and no-ripple prediction values for each cell pair and for each cell type specific interaction. To determine the significance of reciprocal pairs, the proportion of cell pairs with a significant B→A connection was compared between cases where A→B was significantly predictive and A→B was not significantly predictive using a two-sided χ^2^ test of proportions. The significance of predictive coupling for an individual pair was determined by testing whether the predictive filter outputs during co-ripples for that pair were greater than zero using a one-sample one-sided Student’s t-test. To determine the significance of reciprocal triplets, the proportion of significant coupling between a recurrent C→A connection when A→B and B→C are significantly predictive was compared to the proportion of significant C→XX connections, where XX is any other cell outside of the ABC triplet.

The significance of PLV-prediction relationships was determined by bootstrapping analysis. 1000 random selections of pairwise prediction-PLV values were sampled with replacement and fit with a linear model. The relationship was considered significant if 95 percent of the fitted sloped were greater than 0. To determine the significance of prediction-phase relationships, the mean prediction and phase preference for all cell pairs was least-squares fit with a sine curve. The amplitude of the fit sine curve was compared to the amplitude of sine curves fit to data where the pairwise prediction phase values were shuffled 1000 times (Supplementary Figure 7). The mean co-ripple prediction during ripple peaks was compared to ripple troughs for all cell pairs with positive prediction using a paired one sample Student’s t-test.

All fits were approximated with a linear least squares regression, and for fits with R^2^ < 0.3, exponential least squares regressions were instead used if they met R^2^ > 0.3. Fits are only shown for significant linear relationships or well-approximated exponential relationships.

## Acknowledgements

We thank Eran Mukamel,, Mikio Aoi, Charles Dickey, Adam Niese, Dan Soper, Christopher Gonzalez, and Jacob Garrett for their support. This work was supported by NIMH (1RF1MH117155-01, T32 MH020002) and ONR-MURI (N00014-16-1-2829). NINDS (R25NS065743), American Academy of Neurology Clinical Research Training Scholarship, Harvard Catalyst KL2/CMeRIT (DBR). Department of Veterans Affairs (N2864C, A2295R), NINDS (UH2NS095548, U01DC017844), Conquer Paralysis Now 004698, Massachusetts General Hospital-Deane Institute (LRH).

