## Supplementary Figures for "Co-occurring ripple oscillations facilitate neuronal interactions between cortical locations in humans"

### Supplementary Information

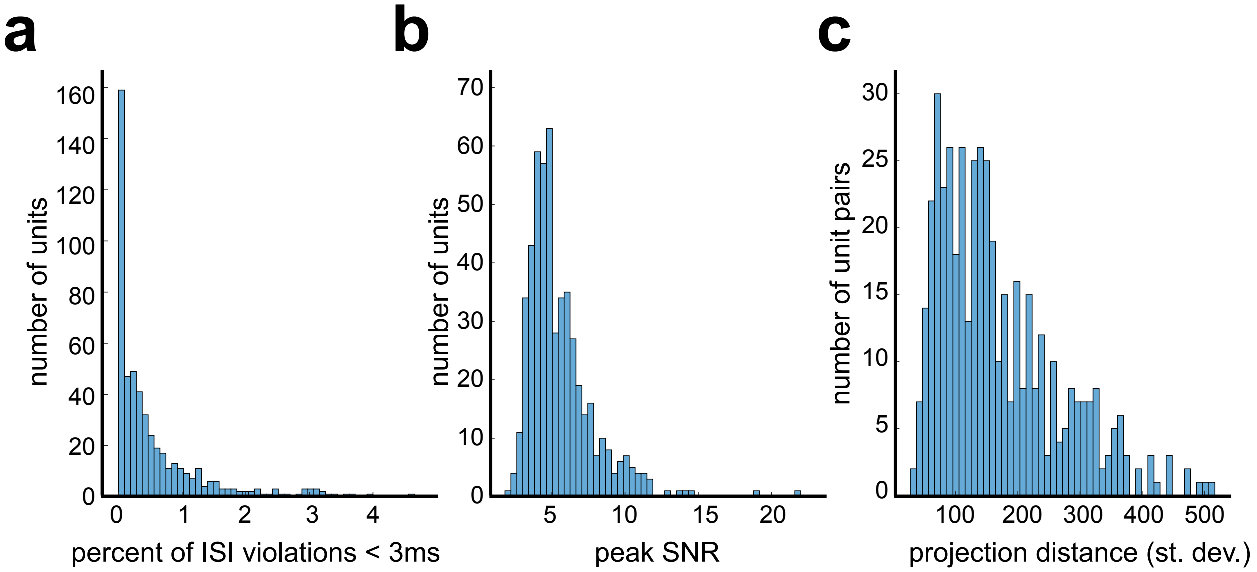


**Supplementary Figure 1. Single unit quality metrics. a** Percent of inter-spike intervals (ISIs) that violate the 3 ms refractory period for each detected single unit. The mean ± st. dev. was 0.55% ± 0.75% across all neurons, indicating low contamination. **b** Signal-to-noise ratio (SNR) at the waveform peak for each detected single unit. The mean ± st. dev. was 5.56 ± 2.26, suggesting that detected units significantly extend beyond the noise floor. **c** Pairwise distance for single units detected on the same Utah array electrode, estimated using the projection test. Units were well separated on the same contact; mean ± st. dev. projection distance was 168.5± 97.

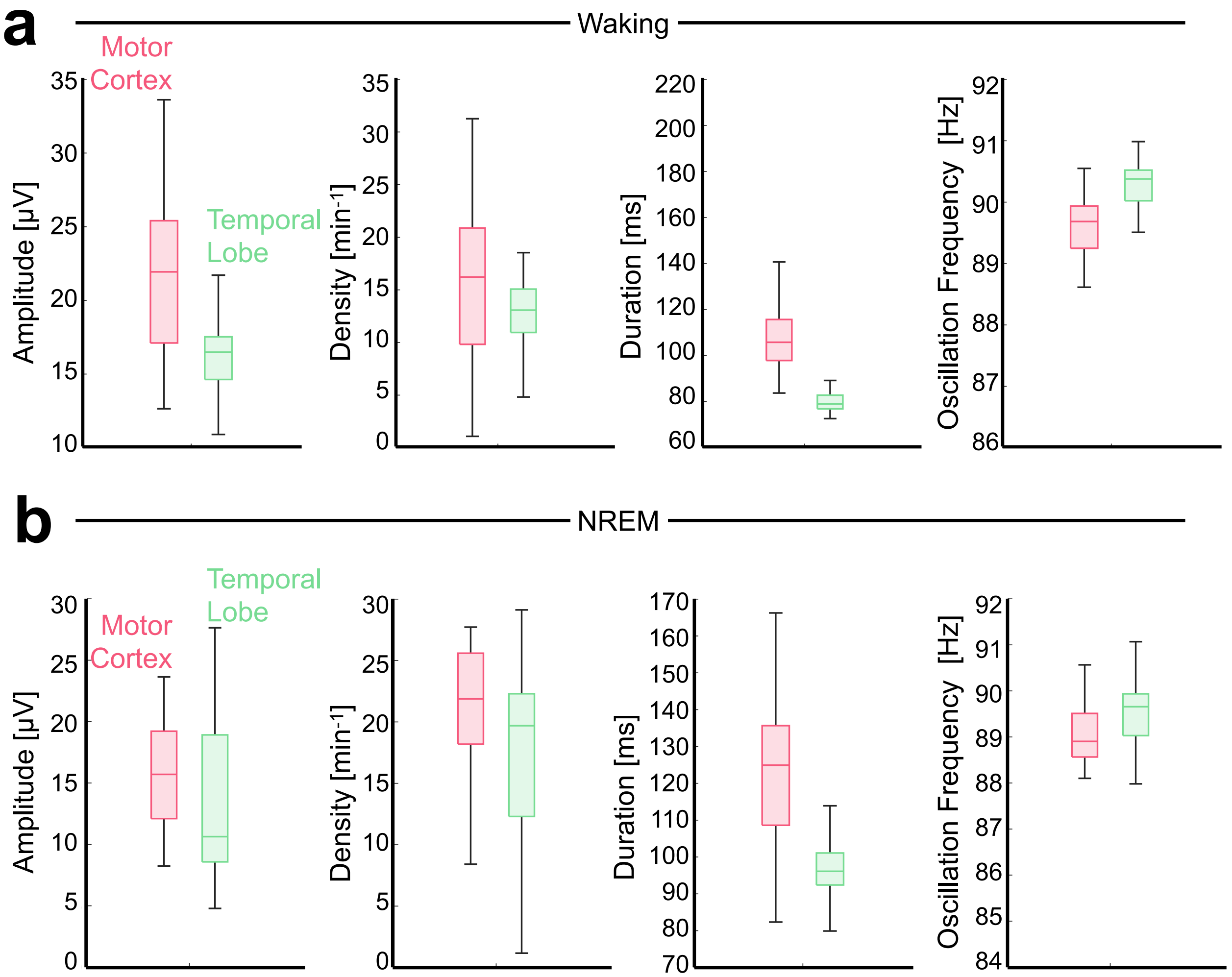


**Supplementary Figure 2. Comparison of ripple characteristics between motor cortex and temporal lobe.** Box plots showing comparison of ripple per-channel amplitude, density, duration, and oscillation frequency during wake (**a**) and NREM (**b**). Horizontal line shows median, box shows upper and lower quartiles, and whiskers show non-outlier maximum and minimum. Patients with arrays implanted in temporal lobe have longstanding epilepsy, while patient with arrays in motor cortex has no known cortical disease.

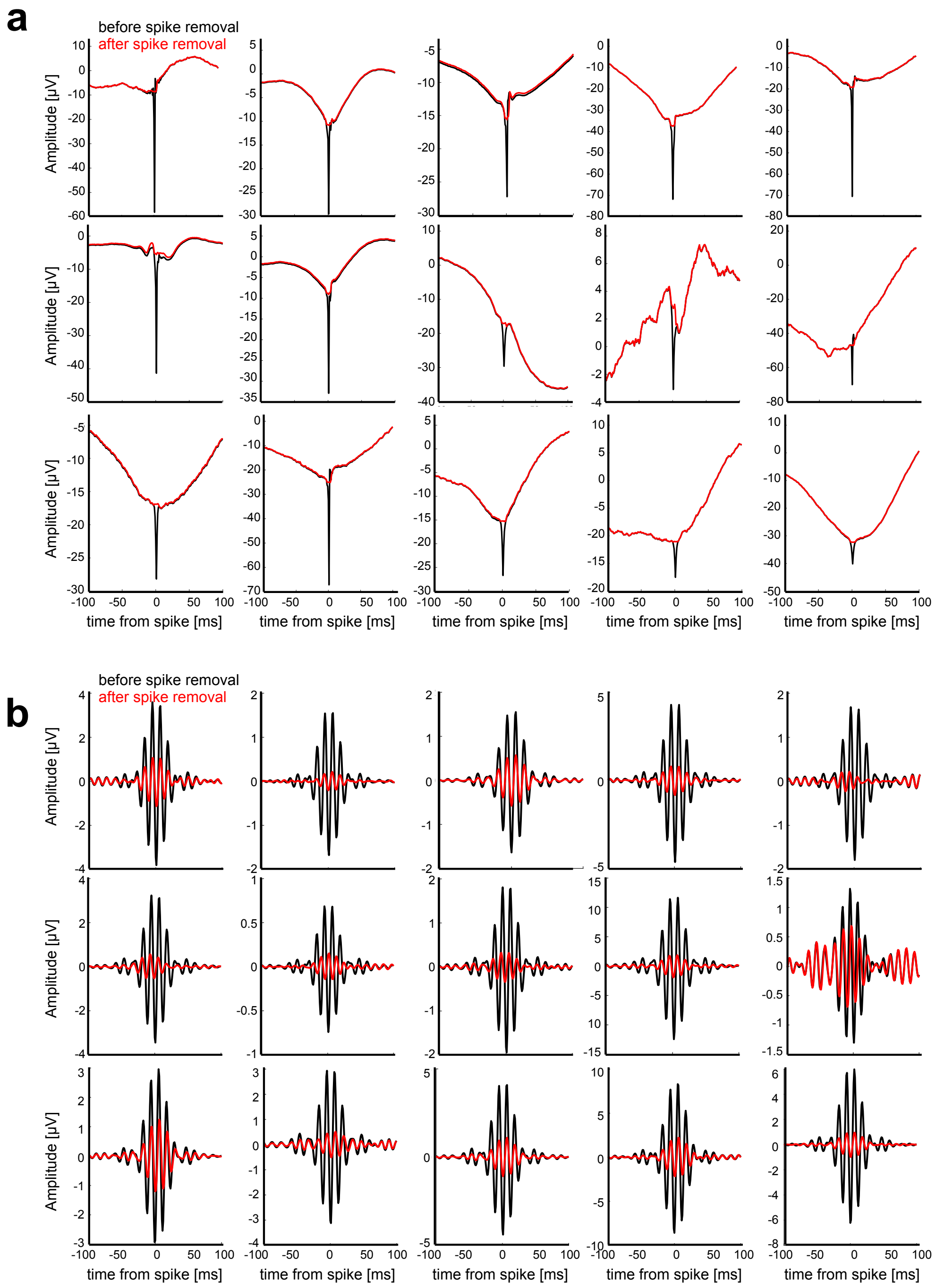


**Supplementary Figure 3. Example of spike removal efficacy in the lowpass LFP. a** Example of spike triggered average (STA) of the broadband LFP before (black) and after (red) removal of the unit waveform from the original high sampling rate data. **b** same as **a**, but showing the STA of the filtered rippleband (70-100 Hz) data. Note that after spike removal, the rippleband filtered data does not exceed 1 µV.

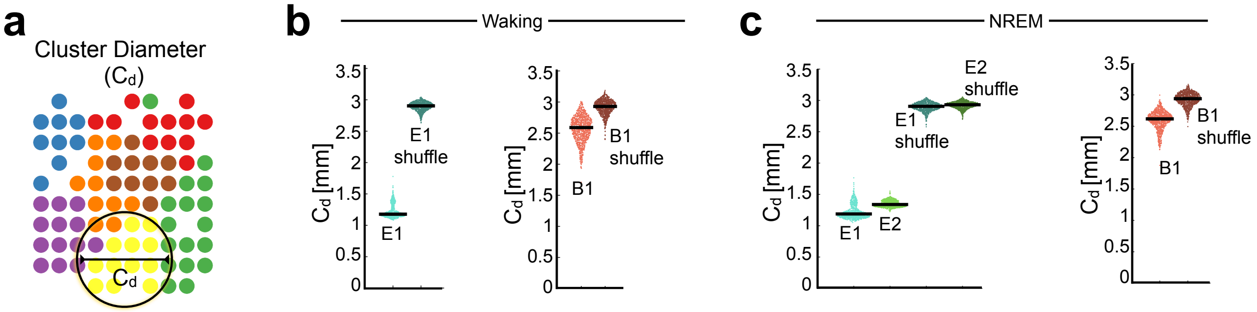


**Supplementary Figure 4. Estimate of phase-locking value cluster size. a** Example of clustering output from nonnegative matrix factorization (NNMF) of the pairwise phase locking value (PLV) matrix for patient E2. Diameter is estimated by computing the radius of gyration for all cluster channel locations compared cluster centroid. **b** Cluster diameter output for multiple iterations of NNMF on waking data, compared to data where the PLV and channel locations are shuffled. **c** Same as **b** but during NREM. Note that the measured cluster diameter in patient.

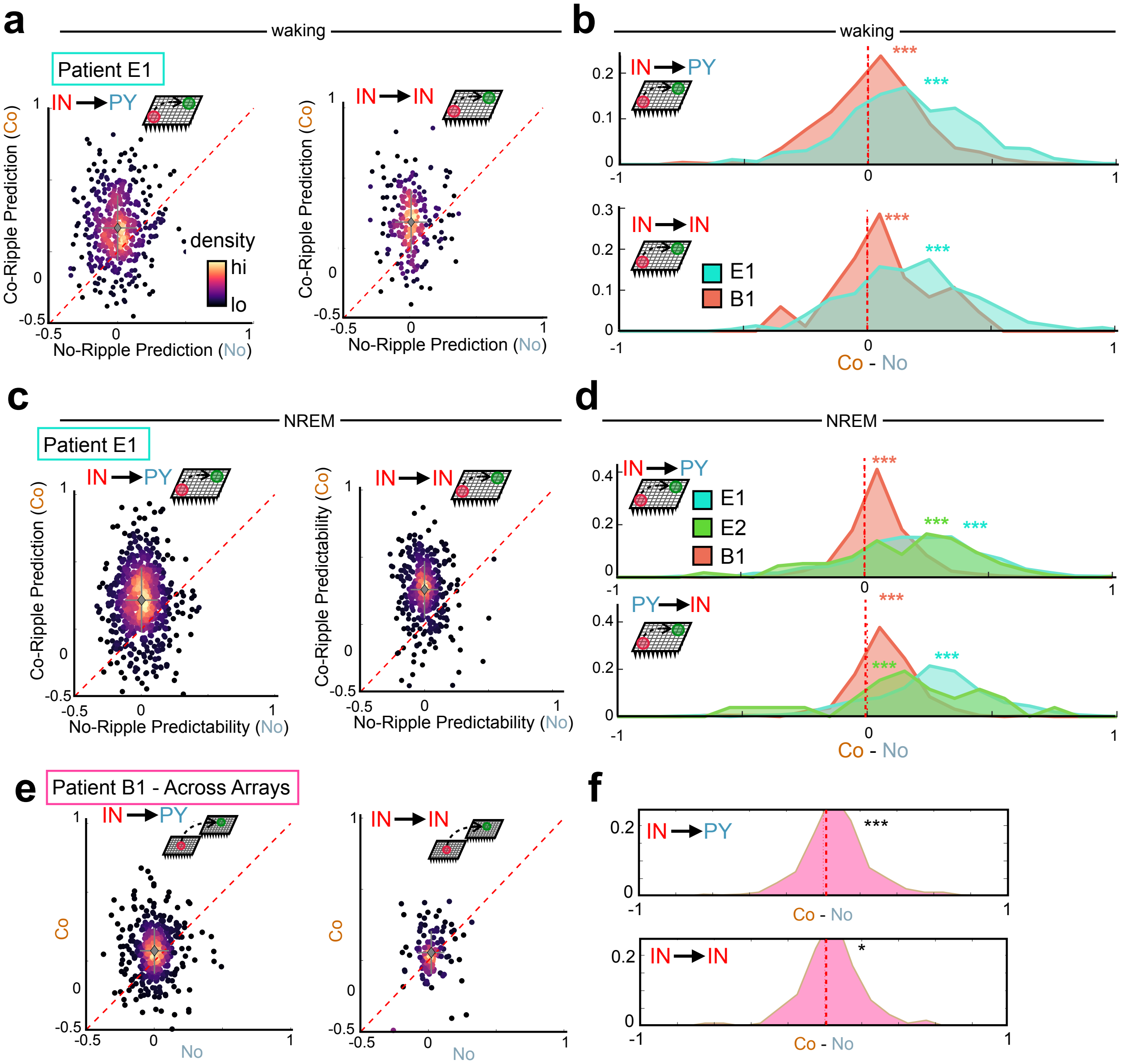


**Supplementary Figure 5. Prediction analysis for IN leading cell pairs. a,b** Same as Fig 4ef, but showing prediction for IN→PY and IN→IN interactions during waking. **c,d** Same as Fig 4gh, but showing prediction for IN→PY and IN→IN interactions during NREM. **e,f** Same as Fig 4i, but showing prediction for IN→PY and IN→IN interactions across both array in patient B1 during NREM. *** $p < 0.001$ * $p < 0.05$ paired two-sided Student's t-test.

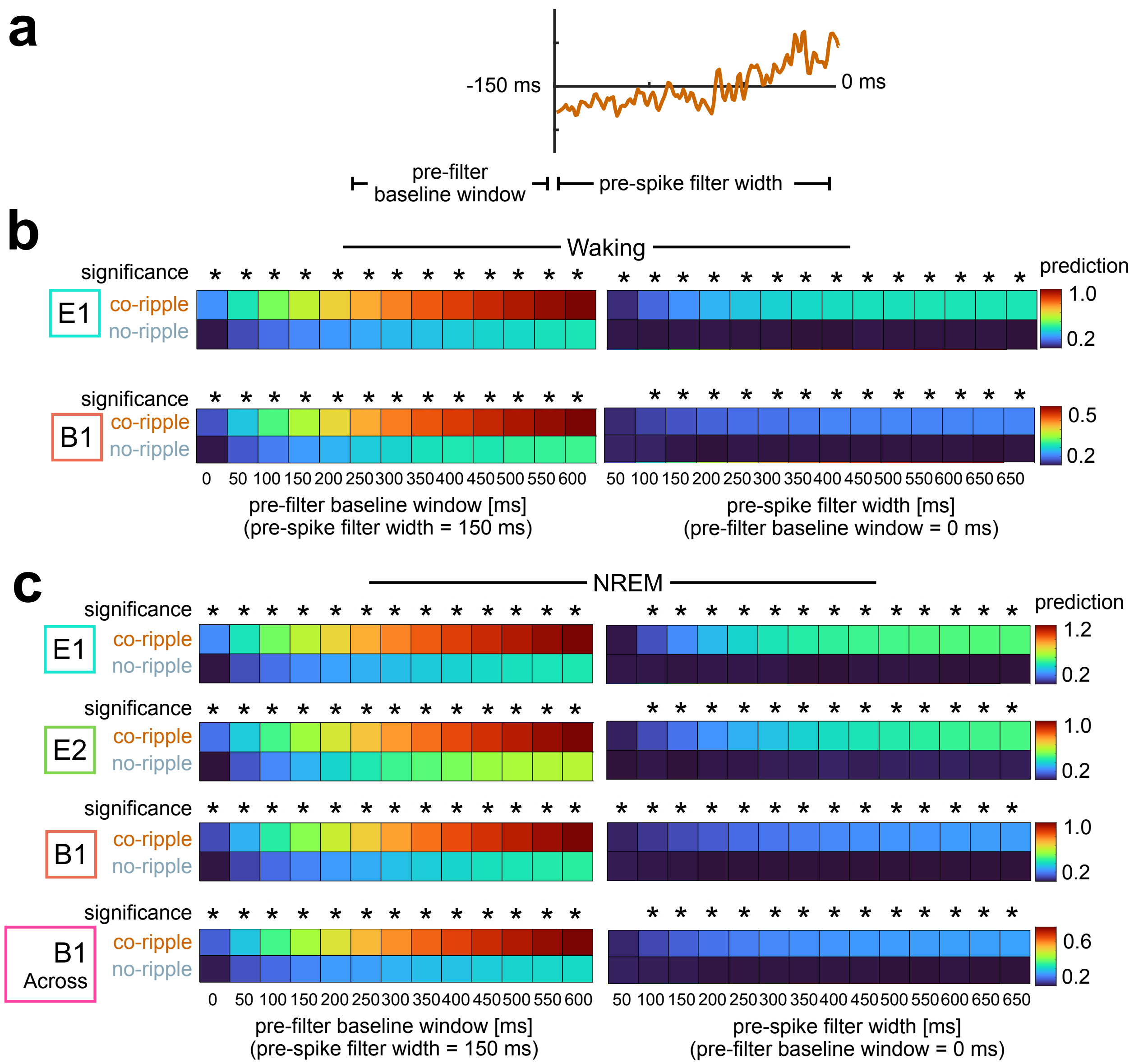


**Supplementary Figure 6. Parameter sweep of the prediction analysis. a** Illustration of parameters explored. Both the filter width and the pre-filter baseline window were explored. **b** Co-ripple and no-ripple results for all parameters explored during waking. Significance between co-ripple and no-ripple values are shown with an asterisk (paired one-sided Student's t-test, $\alpha$ = 0.0001). **c** Same as **b** but for all data in NREM. Parameters chosen for the study, pre-spike filter width: 150 ms, pre-filter baseline window: 0 ms.

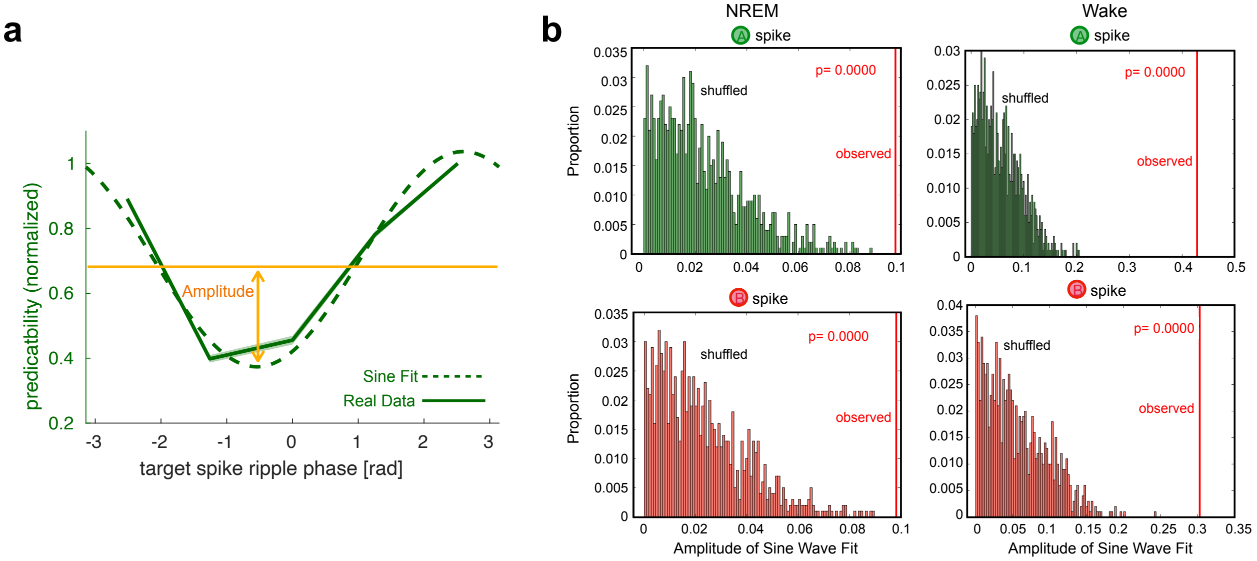


**Supplementary Figure 7. Shuffle analysis of prediction modulation by unit phase preference.** **a** Illustration for how prediction modulation by phase was determined. Modulation index was determined by the amplitude of the sine wave that was fit to the binned prediction-phase relationships. The prediction-phase pairs were shuffled and the amplitude of each resulting sine wave fit were compared to the real data. **b** Shuffle analysis of prediction relationship to target (top) and driver (bottom) neuron phase during NREM. **c** Same as **b**, but during waking.

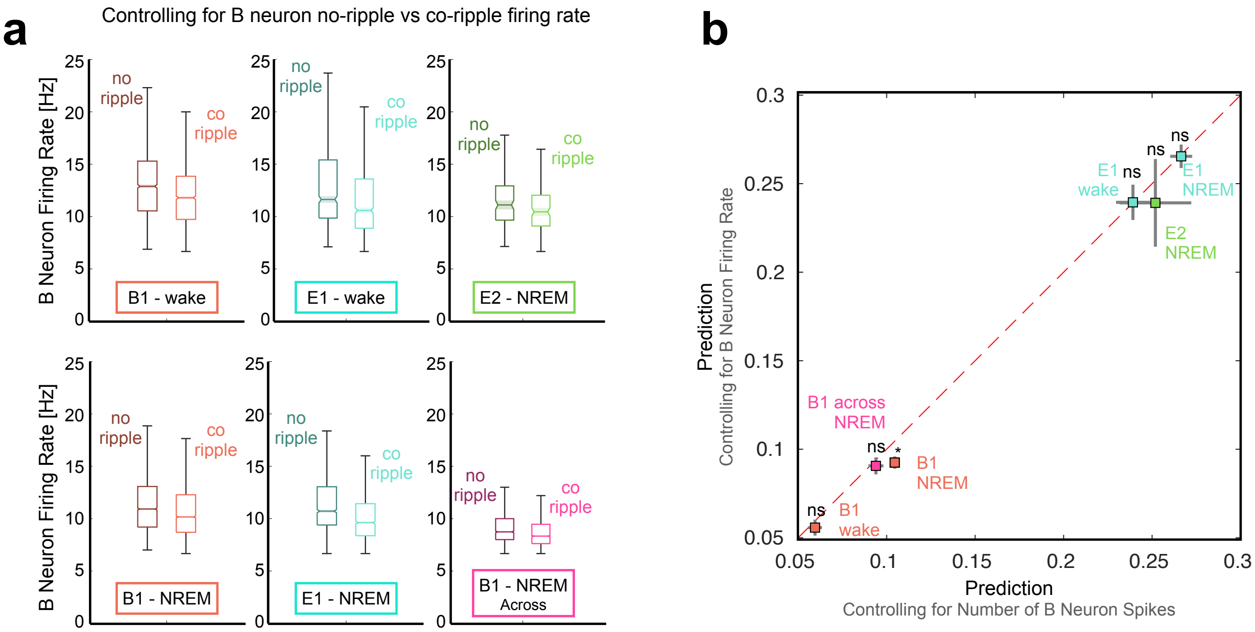


**Supplementary Figure 8. Effect of reducing co-ripple firing rate on prediction.** For each cell pair, trials were removed from the co-ripple condition until the firing rate was below the no-ripple condition. **a** Firing rate for co-ripple and no-ripple conditions for each patient and behavioral state after trials were removed from the co-ripple cases. **b** Comparison between firing rate control method and method presented in study, which is controlled for the number of driver (B) neuron spikes in co- and no-ripples. Red dashed line is x=y. * $p < 0.001$ paired two-sided Student's t-test.

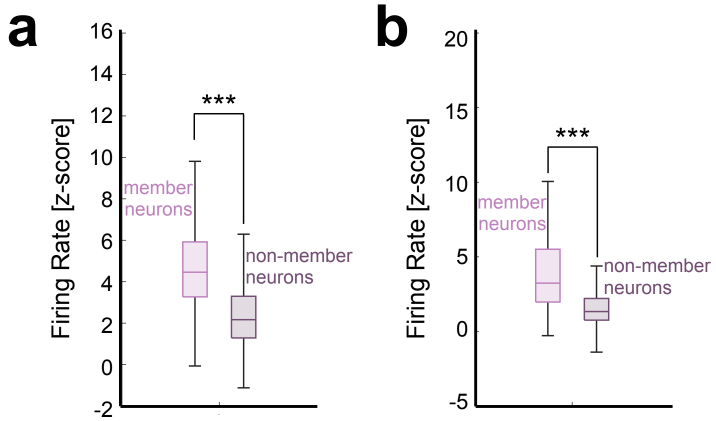


**Supplementary Figure 9. Firing rates during assembly activation.** Z-scored firing rates for member and non-member neurons during assembly activation (strength > 95$^{th}$ percentile). Data is shown for waking (**a**) and NREM (**b**). Horizontal line shows median, box shows upper and lower quartiles, and whiskers show non-outlier maximum and minimum. *** p < 0.001, two-sided rank sum test.

| Waking | | | | | |
|---|---|---|---|---|---|
| **PY↔PY** | | **PY↔IN** | | **IN↔IN** | |
| co-ripple | no-ripple | co-ripple | no-ripple | co-ripple | no-ripple |
| 0.719±0.009 | 0.439±0.006 | 1.08±0.022 | 0.520±0.014 | 2.078±0.083 | 0.703±0.035 |
| **NREM – within array** | | | | | |
| **PY↔PY** | | **PY↔IN** | | **IN↔IN** | |
| co-ripple | no-ripple | co-ripple | no-ripple | co-ripple | no-ripple |
| 0.525±0.006 | 0.160±0.002 | 0.792±0.014 | 0.212±0.005 | 1.710±0.052 | 0.400±0.016 |
| **NREM – across array** | | | | | |
| **PY↔PY** | | **PY↔IN** | | **IN↔IN** | |
| co-ripple | no-ripple | co-ripple | no-ripple | co-ripple | no-ripple |
| 0.348±0.007 | 0.145±0.003 | 0.822±0.037 | 0.322±0.015 | 1.009±0.086 | 0.400±0.037 |

**Supplementary Table 1. Co-fire rates during co-ripple and no-ripple conditions.** Mean ± SEM co-fire rates in Hz for all unit pairwise interactions across waking and NREM. Co-firing is defined as a co-ripple period with a spike in either channel. No-ripple periods were duration matched to each co-ripple event for a given unit pair.
